# Extracellular vesicles facilitate the horizontal transfer of drug resistance and stem-like properties between ovarian tumor cells

**DOI:** 10.64898/2026.01.20.699925

**Authors:** Venkatesh Pooladanda, Rui Xu, Dominique T. Zarrella, Yusuke Matoba, Chisa Shimada, Shaan Kumar, Eugene Kim, Paula Dibenedetto, Xingping Qin, Kristopher A. Sarosiek, Madelyn Krueger, Narela Magrassi, Mansoor Amiji, Maryam Azimi Mohammadabadi, Ursula Winter, Cesar M. Castro, Hyungsoon Im, Raj Kumar, Cheng Wang, Karen D. Cowden Dahl, Kenneth P. Nephew, Oladapo O. Yeku, Lara S. Milane, Bo R. Rueda

## Abstract

Ovarian cancer stem cells (CSCs) can seed recurrent drug-resistant disease. Likewise, non-CSCs can acquire CSC phenotypic properties. How this process is orchestrated is of interest to inform how it might be prevented. We tested the hypothesis that ovarian CSC and/or drug-resistant tumor cells confer stem-like properties via extracellular vesicles (EVs). We focused our investigation on how EVs might mediate EZH2 signaling to promote a phenotypic change in drug-sensitive, non-CSCs. To accomplish this, we utilized paired PARP inhibitor-sensitive and - resistant ovarian cancer (OvCa) cell lines, EZH2 knockdown lines, and patient-derived organoids (PDOs) originating from recurrent high-grade serous OvCa. Small EVs isolated from drug-sensitive, CSC and/or drug-resistant enriched cultures, PARP inhibitor (olaparib) resistant lines, or drug-treated (olaparib or carboplatin) lines were cultured with treatment naïve or sensitive lines for defined time points. The impact of small EV exposure was determined by assessing cell number, metabolic activity, viability, sphere and colony-forming capacity, ALDH activity, DNA damage, and changes in associated signaling pathways. We found that EVs from CSC or drug-resistant enriched cell fractions communicate CSC-like phenotypes to the more sensitive tumor cells via EZH2 canonical and non-canonical signaling pathways, promoting stemness. We conclude that EV-mediated activation of EZH2 signaling represents a targetable mechanism contributing to stemness-associated drug resistance in OvCa.

**Graphical abstract:** 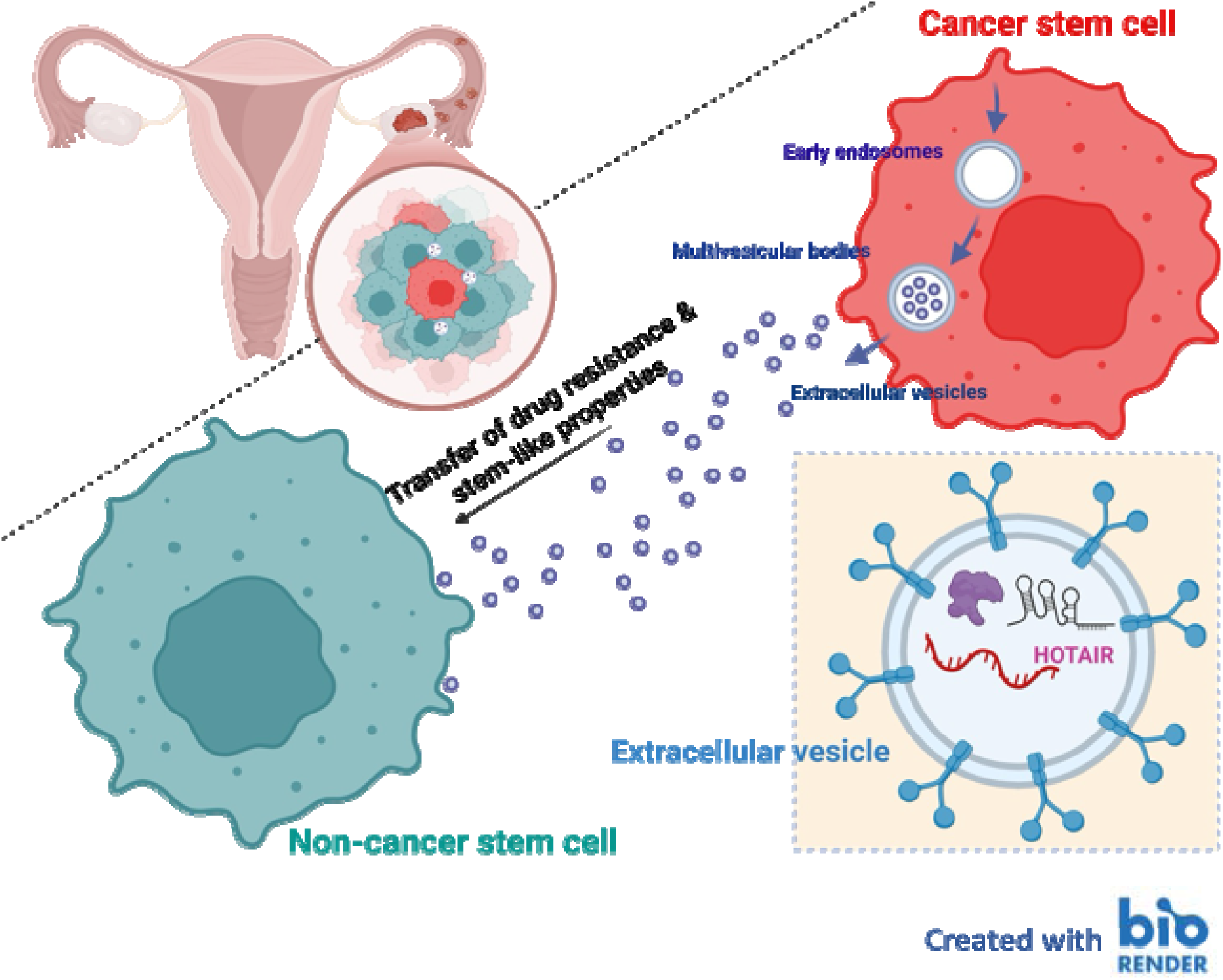

## 1. INTRODUCTION

Ovarian cancer (OvCa) remains one of the most lethal gynecologic cancers in the US, with a 5-year survival rate of about 51% (Siegel et al., 2025). Estimates suggested approximately 20,890 women will be newly diagnosed, while 12,730 women will succumb to the disease in 2025 (Siegel et al., 2025). The high rate of lethality is primarily attributed to the inability to detect early-stage disease and the development of early recurrent, chemoresistant disease (Pinato et al., 2013). The current standard treatment strategy for OvCa includes surgical debulking followed by platinum-based chemotherapy (Armstrong et al., 2006; Ozols et al., 2003) or neoadjuvant therapy with interim cytoreduction followed by additional cycles of chemotherapy. Even with such efforts, patients with high-grade serous OvCa (HGSOC) develop platinum resistance, leading to almost invariably fatal refractory disease with only 43% 5-year overall survival rate for all stages combined (Lokadasan et al., 2016).

The introduction of Poly ADP ribose inhibitors (PARPi) into treatment regimens, which were explicitly designed to promote synthetic lethality in tumors that harbored an alteration in a mediator of the homologous recombination repair pathway (i.e., BRCA I or II) provided some enthusiasm, yet their impact on overall survival has been limited. Patients with platinum-sensitive HGSOC with homologous recombination–deficient (HRD) tumors benefit from PARP inhibitors, but about half of HGSOC patients do not have HRD tumors and thus have very limited treatment options at the chemotherapy-resistant stage (Stiegeler et al., 2024). Those responsive to PARPi often succumb to inherent or acquired resistance mechanisms. Among reported resistance mechanisms, we and others previously demonstrated PARPi’s were much less effective on cancer stem cells (CSC) compared to non-cancer stem cell populations (Bellio et al., 2019a; Bellio et al., 2019b); this enriched CSC populations. Interestingly, the overall increase in CSC numbers following treatment was not concordant with cell proliferation; this difference was further supported by the observation that the bulk of CSC-like cells had an extended G2 phase. This led us to postulate that maybe the non-stem tumor cells exposed to cytotoxics or PARPi’s were displaying some level of plasticity. We refer to plasticity as a cell’s ability to shift between different phenotypical states. The plasticity of tumor cells is not a new concept (Ye and Weinberg, 2015) and has been proposed in OvCa (Wang et al., 2018). Wang et al showed that the more differentiated tumor cells have the potential to shift to a CSC phenotype or vice versa under an appropriate stimulus. Their findings, along with ours, suggest that stress-induced enrichment of CSC is due to conversion of non-CSC to CSC, rather than the expansion of the existing CSC population. Genetic, epigenetic, pharmacologic, and/or microenvironmental changes can trigger the bidirectional interconversion between stem and non-stem-like states (van Neerven et al., 2016). The question remains how this might be mediated in OvCa.

EVs are cell-derived vesicles contained by a lipid bilayer. Small EVs, often referred to as exosomes, are formed within the cell and released via multivesicular vesicles, whereas ectosomes (a.k.a. microparticles) develop from the cell membrane. Despite their small sizes, EVs carry nucleotides (RNA, DNA, microRNA, long noncoding RNAs (lncRNA), proteins, and lipids to communicate with other cells within and outside the cellular microenvironment (Wessler and Meisner-Kober, 2025). Functionally, EVs promote a range of biological functions, including cell differentiation, proliferation, angiogenesis and/or regulate immune response and promote drug resistance in tumor cells.

EVs derived from stromal cells in the tumor microenvironment have been shown to support the maintenance and/or enrichment of the CSC population (Brown and James, 2021). Since we observed shifts in phenotype within a fixed cell population of OvCa cells, we explored the potential for EVs to drive the exchange of CSC-like phenotypes between ovarian tumor cells and whether the shift in phenotype was mediated, at least in part, by epigenetic modification.

Enhancer of Zeste Homologue 2 (EZH2), a histone methyl transferase, is a key catalytic component of the polycomb repressor complex 2 (PRC2), which is most well-known for its epigenetic modifications by catalyzing the trimethylation of lysine 27 on histone 3 (H3K27me3). Dysregulated EZH2 expression is implicated in several cancers, including OvCa, where EZH2 is frequently overexpressed and associated with poor prognosis, increased tumor aggressiveness, resistance to therapy and promoting stemness (Jones et al., 2018; Reid et al., 2021; Rizzo et al., 2011). Our objective was to demonstrate that EVs mediate phenotypic plasticity or dedifferentiation in OvCa cells by altering the epigenome through increased expression and activity of EZH2.

## 2. Experimental Section

For a more detailed methodology, see the Supplemental Materials section.

### Cell lines and culture conditions

Human epithelial OvCa cell lines were obtained from multiple sources. A2780 (RRID: CVCL_0134) cells were kindly provided by Dr. Karen Cowden-Dahl (Kabara Cancer Research Institute, La Crosse, WI). UWB1.289 mut cells (RRID: CVCL_B079) were purchased from the American Type Culture Collection (ATCC, Manassas, VA). OVCAR4 cells (RRID: CVCL_1627) were obtained from Sigma-Aldrich (Burlington, MA). The PEO1 cell line (RRID: CVCL_2686) was generously provided by Dr. Kevin Elias (Brigham and Women’s Hospital, Boston, MA). A2780 and OVCAR4 cells were maintained in RPMI-1640 medium (Cat #11,875,093, Gibco, Gaithersburg, MD) supplemented with 10% fetal bovine serum (FBS) (Cat #26,140,079, Thermo Fisher Scientific, Waltham, MA) and 1% penicillin-streptomycin (Pen-Strep) (Cat # 15,070,063, Thermo Fisher Scientific). UWB1.289 mut cells were cultured in a 1:1 mixture of RPMI-1640 and MEGM (Mammary Epithelial Growth Medium) (Cat #CC-3150, Lonza, Cambridge, MA) supplemented with 10% FBS and 1% Pen-Strep. PEO1 cells were maintained in RPMI-1640 medium supplemented with 10% FBS and 1% Pen-Strep. For the EV experiments, cells were cultured and/or treated in exosome-free serum medium (Cat #A2720801, Thermo Fisher Scientific). All cell lines were cultured under standard conditions in a humidified incubator at 37°C with 5% CO_2_. The cell lines were authenticated using STR profiling (Labcorp, Burlington, NC) and confirmed to be negative for mycoplasma using a MycoAlert Mycoplasma Detection Kit (Cat# LT07-318, Lonza) before any experiments were performed. No contamination was detected during the study period.

### HGSOC patient-derived organoid (PDO) cultures

17-121 PDO established from a recurrent patient with BRCA mutation and exhibited olaparib resistance and VCRB330 PDO established from the recurrent patient with multiple treatments (carboplatin, paclitaxel, cisplatin and gemcitabine) before sample collection. These PDO cultures were established and maintained as previously described (Hill et al., 2018). Organoids were monitored for growth and passaged as needed based on their growth rate.

### Assessment of ALDH activity

Cells were seeded in a 6-well plate, and the percentage of ALDH activity was determined using the ALDEFLUOR Kit (Cat #01700, StemCell Technologies, Cambridge, MA) and flow cytometry (Beckman Coulter Gallios, Brea, CA) according to the manufacturer’s guidelines. Ovarian cancer cell populations that display a high percentage of ALDH activity are enriched for cells that display CSCs properties relative to the corresponding controls displaying a low percentage of ALDH activity (Yokoyama et al., 2016).

### Generation of EZH2 knockdown UWB 1.289 mut cell lines

Lentiviral particles were produced by co-transfecting HEK293T cells procured from ATCC with the EZH2 Human shRNA Plasmid (Cat #TL304713, OriGene, Rockville, MD) and packaging plasmids, followed by collection, filtration, and concentration of the viral supernatant. UWB 1.289 cells were transduced with the lentiviral particles in the presence of 5 µg/mL polybrene, and after 24 hours, the medium was replaced. Transduced cells were selected with 1 µg/mL puromycin for 7 days, expanded, and maintained in medium with 1 µg/mL puromycin. GFP-positive cells were sorted using flow cytometry, and sorted cells were cultured in culture medium with 1 µg/mL puromycin. EZH2 knockdown efficiency was validated by western blotting.

### Western blot analysis

EZH2, H3K27me3, and phosphorylated CHK1 (Ser296 and Ser345) levels were determined using standard western blot techniques as previously described (Bellio et al., 2019a; Cardenas et al., 2016; Matoba et al., 2024). Specific details related to antibodies and dilutions are provided in Supplementary Materials. Membranes were incubated overnight at 4°C with primary antibodies specific to EZH2 (1:1000), H3K27me3 (1:1000), pCHK1 Ser296 (1:1000), and pCHK1 Ser345 (1:1000). Antibody details are provided in the supplementary material. Detection of protein bands was performed using an enhanced chemiluminescence (ECL) substrate and captured on ChemiDoc Imaging System (Bio-Rad). Protein levels were quantified using ImageJ software, and β-Actin and Histone H3 served as loading controls to normalize for differences in protein loading.

### Spheroid forming assay

Spheroids were generated as previously described (Bellio et al., 2019a; Matoba et al., 2024). Briefly, cells were seeded in ultra-low attachment 6-well plate (Cat #7007, Corning, Glendale, AZ) at a density of 20,000 cells per well in serum-free medium supplemented with growth factors (e.g., EGF, bFGF, and B-27™ supplement) and incubated at 37°C in 5% CO_2_. Spheroid formation was monitored over 13 days, with spheroid medium replenished every 3 days. At the end, spheroids and media were collected, allowed to settle, and resuspended in phenol red-free medium. Spheroid number and size were quantified using ImageJ software, where spheroids were defined as objects with a pixel size greater than 50 pixels.

### Colony-forming assay

The colony-forming assay was performed as we previously described (Matoba et al., 2024). Briefly, cells were seeded at a density of 500 cells per well in 6-well plates. The cells were maintained under standard cell culture conditions and incubated for 14 days. Following the incubation period, the colonies were fixed with methanol and stained with crystal violet. Images of the stained colonies were captured, and the number of colonies was quantified using ImageJ analysis software.

### Co-culture studies

Cells were co-cultured using 6-well transwell plates with a pore size of 400 nm to facilitate paracrine signaling between the two cell populations while preventing direct cell-cell contact. In this setup, 1 × 10 cells were seeded in 1 mL of appropriate culture media in the upper chamber, while 0.5 × 10 cells were seeded in 2 mL of appropriate culture media in the lower chamber. The co-culture system was maintained for 72 hours under standard cell culture conditions (37°C, 5% CO_2_, and 95% humidity). After the incubation period, cells from the lower chamber were carefully collected and analyzed for ALDH activity as described earlier.

### Generation of olaparib-resistant OvCa cell lines

Olaparib-resistant UWB1.289 mut (BRCA1 null) and PEO1(BRCA2 mutant) cell lines were developed by continuous exposure to increasing concentrations of olaparib (Cat #S1060, Selleckchem, Houston, TX) (0.5 to 50 µM) over a period of 3 months. Resistant cell populations were maintained in the presence of 10 µM olaparib to sustain resistance. Resistance was confirmed by performing MTT cell viability assays and determining the IC_50_ values in comparison to parental drug-sensitive cell lines, which were exposed to DMSO as a control.

### MTT and cell counts

The metabolic activity of cells was assessed using an MTT (3-(4,5-dimethylthiazol-2-yl)-2,5-diphenyltetrazolium bromide) assay as previously described (Matoba et al., 2024). Briefly, cells were seeded in a 96-well plate at a density of 1,000 cells per well. The cells were then treated with either EVs, olaparib, or a combination of both. After the treatment period, the medium was replaced with fresh medium containing 0.5 mg/mL MTT reagent (Cat # M6494, Thermo Fisher Scientific), and the cells were incubated at 37°C in a 5% CO_2_ atmosphere for 4 hours. Following incubation, the supernatant was carefully removed, and the formazan crystals formed were dissolved in dimethyl sulfoxide (DMSO). Absorbance was measured at 570 nm using a microplate reader (SpectraMax, Molecular Devices, San Jose, CA). Viable cells were counted by trypan blue (Cat #15250061, Thermo Fisher Scientific) exclusion. Briefly, cells were trypsinized and washed with PBS and mixed with a 1:1 ratio with 0.4% of trypan blue stain, and viable cells were counted by TC20 Automated Cell Counter (Bio-Rad, Hercules, CA).

### Cell viability and apoptosis assay

To assess cell viability and apoptosis, we utilized the FITC or APC Annexin V Apoptosis Detection Kit with 7-AAD (Cat #640922 and #640930, BioLegend) as previously described (Matoba et al., 2024). Flow cytometric analysis allowed for quantification of the populations, where viable cells were negative for both markers, apoptotic cells were Annexin V-positive, and necrotic cells were 7-AAD-positive but Annexin V-negative.

### EV isolation

EVs were isolated from culture medium following a series of centrifugation and filtration steps. The cell culture medium was first centrifuged at 200 × g for 5 minutes to remove debris, followed by 2,000 × g for 10 minutes at 4°C to eliminate cells. The supernatant was then filtered through a 0.22 µm sterile syringe filter pre-wetted with Dulbecco’s Phosphate Buffered Saline (DPBS) (Cat #14190144, Thermo Fisher Scientific). The filtered supernatant was concentrated using a 100 kDa MWCO centrifugal filter unit (Amicon® Ultra-15 Centrifugal Filter Unit; Cat #UFC910024, MilliporeSigma, St. Louis, MO) at 3,000 × g for 30 minutes at 4°C (Shu et al., 2021). The resulting retentate was washed three times with sterile DPBS to remove the residual medium, and the concentrated EVs were collected. The EVs were further purified and concentrated by ultracentrifugation using a Beckman Coulter Optima MAX-XP Ultracentrifuge with a TLA-55 rotor (Cat #3574478, Beckman Coulter) at 100,000 × g (55K RPM) for 1.5 hours at 4°C (Coughlan et al., 2020). The final EV pellet was resuspended in sterile DPBS, aliquoted (∼200 µL), and stored at -80°C.

### Nanoparticle tracking analysis (NTA)

To obtain small EVs, the size distribution and concentration (number of particles) of EVs were determined using Nanoparticle Tracking Analysis (NTA) using a Nanosight LM10 (Malvern, Framingham, MA). The small EVs were isolated and diluted in PBS to achieve an optimal concentration for experiments (**Table S1**).

### EV characterization by flow cytometry

For EV flow cytometry analysis, 2 µL of beads (Cat #A37304, Thermo Fisher Scientific) were mixed with 5 µL of the EV sample, followed by the addition of 75 µL of cell staining buffer. The mixture was incubated for 15 minutes at room temperature with gentle mixing. A control sample without EVs was also prepared. Fc receptor (FcR) blocking reagent (20 µL) (Cat# 130-059-901, Miltenyi Biotec, Charlestown, MA) was added to each tube, and the samples were incubated on ice for 10 minutes. Following the addition of 500 µL of cell staining buffer, the samples were centrifuged at 3,000 × g for 4 minutes. The supernatant was discarded, and the pellets were resuspended in 200 µL of 100 mM glycine in PBS, followed by a 30-minute incubation. The samples were then centrifuged at 3,500 × g for 4 minutes, and the supernatant was removed. The pellets were resuspended in 300 µL of cell staining buffer. Flow cytometry antibodies (FITC-CD9, PE-CD63, and APC-CD81, Biolegend) were added at a final concentration of 1 µL per 100 µL of sample and incubated for 1 hour at room temperature. After incubation, 500 µL of cell staining buffer was added, and the samples were centrifuged at 3,500 × g for 4 minutes, with the supernatant discarded. This washing step was repeated twice to remove unbound antibodies. Flow cytometry analysis was then performed to characterize the EV surface markers. EV protein concentration was determined by microBCA assay kit as per manufacturer guidelines (Thermo Fisher Scientific, US). Calnexin protein expression was tested by western blotting as described (Gomes et al., 2015) using anti-calnexin (Cell Signaling Technologies), which acts as a negative control for EV analysis. The size and purity of isolated EVs were further confirmed by flow cytometric analysis using the EV-specific markers CD9, CD63, and CD81. The absence of the negative EV marker calnexin was validated by immunoblotting (**Table S2 and Figure S1-3**). Isolated EVs were imaged by transmission electron microscopy (TEM) (**Figure S4**).

### Establishment of A2780 ALDH+ spheroids

A2780 cells were cultured to confluence and harvested for spheroid formation. The cells were first labeled with an ALDH substrate using an ALDEFLUOR™ kit (Cat #01700, StemCell Technologies, Cambridge, MA) according to the manufacturer’s instructions. ALDH+ cells were isolated by fluorescence-activated cell sorting (FACS) (BD FACSAria™, Franklin Lakes, NJ) based on their positive staining for ALDH activity. The sorted ALDH+ A2780 cells were seeded at a density of 20,000 cells per well in ultra-low attachment 6-well plates in serum-free medium supplemented with growth factors to promote spheroid formation. The plates were incubated at 37°C in a humidified incubator with 5% CO_2_, and spheroid formation was monitored over a period of 10–14 days.

### Comet assay

Assessment of DNA damage response in human OvCa cells was performed as we previously described (Bellio et al., 2019a). Cells were seeded in a 6-well plate at an appropriate density. The next day, the cells were primed with EVs derived from olaparib-resistant cells or control cells. On the subsequent day, the cells were treated with 10 μM olaparib or DMSO, in combination with EVs. EVs were replenished daily for a total incubation period of 72 hours. At the end of the incubation, cells were scraped, and 1 × 10 cells were collected for the comet assay, following the manufacturer’s instructions from the Comet Assay Kit (Cat #7100, Cell BioLabs, San Diego, CA). Briefly, cells were resuspended, embedded in agarose on microscope slides, and subjected to electrophoresis. DNA damage was assessed by the appearance of comet tails, indicative of DNA strand breaks. Images were captured using a fluorescence microscope at 200× magnification.

### RT-qPCR for HOTAIR

Total RNA was extracted from the EVs of olaparib-sensitive and resistant cell lines using the RNA extraction kit (Thermo Fisher) according to the manufacturer’s protocol. cDNA was synthesized from total RNA using the Maxima First Strand cDNA Synthesis Kit (Thermo Fisher) following standard procedures. Quantitative PCR was performed to measure HOTAIR expression using the SsoAdvanced Universal SYBR Green Supermix (Bio-Rad). The specific primers for HOTAIR were: forward 5′-CAGTGGGGAACTCTGACTCG-3′ and reverse 5′-GTGCCTGGTGCTCTCTTACC-3′, with GAPDH primers used for normalization: forward 5′-GTCAACGGATTTGGTCTGTATT-3′ and reverse 5′-AGTCTTCTGGGTGGCAGTGAT-3′ (Kogo et al., 2011). The PCR reaction was performed on a CFX96™ Real-Time PCR System (Bio-Rad). Relative expression levels of HOTAIR were calculated using the ΔΔCt method, normalizing to GAPDH.

### BH3 profiling assay

Based on preliminary EV exposure curves, UWB-OlaSen cells were treated with EVs isolated from UWB-OlaRes at a concentration of 2.5 µg/mL for 24, 48, and 72 hours, with control EVs (from UWB-OlaSen cells) included for comparison. Following the treatment period, BH3 profiling was performed to assess mitochondrial apoptotic priming and dependencies. The assay was conducted as described in a previously published protocol (Fraser et al., 2019) with slight modifications. Briefly, treated cells were harvested and resuspended in Mannitol Experimental Buffer 2 (MEB2), consisting of 10 mM HEPES (pH 7.5), 150 mM mannitol, 150 mM KCl, 1 mM EGTA, 1 mM EDTA, 0.1% BSA, and 5 mM succinate, then added to prepared plates containing the indicated peptide conditions and 0.001% digitonin. After incubation for 60 minutes at 28°C, cells were fixed with 8% paraformaldehyde for 15 minutes and neutralized with Neutralizing (N2) buffer (1.7 M Tris base, 1.25 M glycine, pH 9.1). Cells were stained overnight with anti-Cytochrome c-Alexa Fluor 647 (1:2000, clone 6H2.B4, Biolegend) and DAPI (1:1000, Abcam) in an intracellular stain buffer (0.2% Tween20, 1% BSA). Cytochrome c release was analyzed by flow cytometry (Attune NxT) (Thermo Fisher Scientific) the following day. Data were analyzed to evaluate differential responses to EV treatment at the specified time points.

### Mitochondrial network analysis

UWB1.289 mut cells were cultured in complete medium at 37°C with 5% CO and seeded at a density of 75,000 cells per well in 35 mm glass-bottom Ibidi dishes. After 24 hours, the medium was replaced with exosome-depleted media, and cells were incubated for an additional 24 hours. Baseline mitochondrial dynamics were assessed in both UWB1.289 mut OlaSen and OlaRes cells. To evaluate the impact of EVs, OlaRes cells were treated with EVs derived from OlaRes cells (2.5 µg/ml), while EVs from OlaSen cells served as controls. Additionally, proMFN2 peptide-loaded liposomes (10 µM) were used as a positive control. Mitofusin 2 (MFN2) mediates mitochondrial network fusion at the level of the outer mitochondrial membrane, and the proMFN2 peptide is a validated peptide that increases mitochondrial networks (Franco et al., 2016). Following 24 hours of treatment, cells were stained with 50 nM Mitotracker Green FM for 15 minutes at 37°C. Live-cell imaging was performed using a Zeiss LSM 880 Confocal Fluorescence Microscope equipped with a 63X objective lens. Mitochondrial dynamics were quantified, including the number of individual mitochondria, the number of mitochondrial networks, and the size (branch length and number of branches) of networks using ImageJ Fiji and Mitochondrial Network Analysis software (Valente et al., 2017). In Fiji, the fluorescent channel of each image was processed by unsharpening the mask, enhancing local contrast, converting to binary (for the mitochondrial footprint measurement), skeletonizing, and analyzing the 2D/3D skeleton with Mitochondrial Network Analysis software. Statistical comparisons were conducted to evaluate mitochondrial network changes across experimental groups.

### Statistical analysis

Data were analyzed using appropriate statistical tests to determine significant differences between experimental groups. Student’s *t*-test was used to compare the means of two groups, while one-way and two-way analysis of variance (ANOVA) were employed to compare means across multiple groups. For ANOVA, post-hoc Tukey’s multiple comparisons test was performed to identify specific pairwise differences between groups. All statistical analyses were performed using GraphPad Prism (GraphPad Software, La Jolla, CA), and results were considered statistically significant at a p-value < 0.05. Data are presented as mean ± standard error of the mean (SEM), as specified in the figure legends.

## 3) RESULTS

### 3.1. Pharmacologic inhibition and knockdown of EZH2 via shRNA negatively impact the ovarian CSC population and inhibit PARPi-induced increase in CSCs

Others have demonstrated that pharmacological inhibition and knockdown of EZH2 via shRNA negatively impact the CSC population in various OvCa cell lines (Wen et al., 2021; Zong et al., 2020). To confirm that EZH2 mediates CSC levels and activity in our preclinical models, as well as its impact on PARPi-induced enrichment of CSCs, we employed two approaches. First, we used GSK126, a small molecule inhibitor designed to disrupt the catalytic activity of EZH2.

Second, we knocked down EZH2 using a shRNA-based strategy. While several markers are used for the identification, isolation, and enrichment of ovarian CSCs, ALDH activity is one of the more widely accepted markers, aside from *in vitro* and *in vivo* functional assays. As anticipated based on our previous study (Bellio et al., 2019a), treatment with olaparib alone increased ALDH activity in UWB1.289 mut, PEO1, A2780, and OVCAR4, and co-treatment with GSK126 inhibited the olaparib-induced increase in ALDH activity across all OvCa cell lines tested (**Figure S5A-D**). Flow cytometric analysis revealed that GSK126 effectively suppressed the drug-induced expansion of the stem-like population in 3 of the 4 cell lines, suggesting that EZH2 inhibition can mitigate the enhancement of ALDH activity, a common marker of CSC, induced by PARP inhibition.

To further confirm the functional role of EZH2 in OvCa cells, we knocked down EZH2 in UWB1.289 mut cells using an EZH2 human shRNA-based strategy. Western blot analysis confirmed a reduction of EZH2 expression in both EZH2 shRNA clone 1 (C1) and clone 2 (C2) compared to scramble shRNA-expressing cells (**Figure S6A**). This reduction in EZH2 was mirrored by a decrease in H3K27ME3 (**Figure S6A**), supporting that EZH2 was driving canonical histone modifications. Reduction of EZH2 expression slowed cell proliferation, as determined by cell counts (**Figure S6B**) and metabolic activity using an MTT assay (**Figure S6C**). Clonogenic assays revealed that EZH2 knockdown reduced colony-forming ability (**Figure S6D, E**). EZH2 knockdown also resulted in a reduced spheroid-forming capacity. Of those colonies that formed from EZH2 knockdown cells from C1, they were smaller in size relative to their controls, but the size of the spheres from C2 was not different when compared to scramble shRNA-expressing cells (**Figure S6F–H**). Percent ALDH activity was reduced in EZH2 knockdown lines compared to the scrambled control cell line (**Figure S6I, J**). Together, these findings demonstrate that the reduction of EZH2 interferes with OvCa cell proliferation, colony and sphere-forming capacity, and ALDH activity, further confirming EZH2’s influence on stem-like activity in OvCa.

### 3.2. Assessing whether indirect cell-cell communication between cell pools is sufficient to enhance stemness properties

To begin to investigate the contribution of cell-cell communication in regulating stemness among cell lines that have different baseline levels of CSCs and non-CSCs, we used a transwell co-culture system to examine whether cells with known ALDH activity could impact ALDH activity in cells that displayed lower baseline levels. To begin, we used the A2780 OvCa cell line, which others and we have shown to have relatively higher baseline levels of ALDH activity (9 to 47%) (Mei et al., 2025; Silva et al., 2011; Uddin et al., 2020) compared to UWB1.289 cells (<3%) (Bellio et al., 2019a). A2780 cells were cultured in the upper chamber, which was separated from the lower chamber by a 0.4 μm porous barrier, which was seeded with UWB1.289 mut cells. Control co-cultures included A2780 cells in both upper and lower chambers, as well as UWB1.289 mut cells in both (**Figure S7A**). After 72 hours, the indirect interaction between A2780 and UWB1.289 lines increased ALDH activity in the UWB1.289 cells (**Figure S7B, C**). These findings support our hypothesis that some signal(s) could be exchanged that could influence cells to display a phenotypic marker linked to CSC and support the concept that EVs may be involved in facilitating this conversion.

Given our focus on CSC enrichment in PARPi-treated OvCa and the potential for EVs to mediate, in part, this enrichment, we generated olaparib-resistant (OlaRes) variants of BRCA1-mutant UWB1.289 (**Figure S8A**) and BRCA2-mutant PEO1 (**Figure S8B**) cell lines. The resistant lines exhibited higher IC_50_ values compared to their olaparib-sensitive (OlaSen) counterparts. Notably, OlaRes cells displayed elevated ALDH activity relative to OlaSen cells in both UWB1.289 mut (**Figure S8C**) and PEO1 (**Figure S8D**) lines. Additionally, sphere-forming assays revealed a marked increase in the number of spheroids formed by OlaRes cells compared to OlaSen cells, although spheroid size remained unchanged in both UWB1.289 (**Figure S8E**) and PEO1 (**Figure S8F**) lines. Next, we again employed our co-culture model (**Figure 1A**) to assess the influence of cells with high ALDH activity on those with low ALDH activity. In both UWB1.289 (**Figure 1B**) and PEO1 (**Figure 1C**) lines, co-culture with their corresponding OlaRes cells increased ALDH activity in OlaSen cells. These data further support the idea that at least ALDH activity, a marker for stemness, can be influenced via indirect communication.

**Figure 1:**
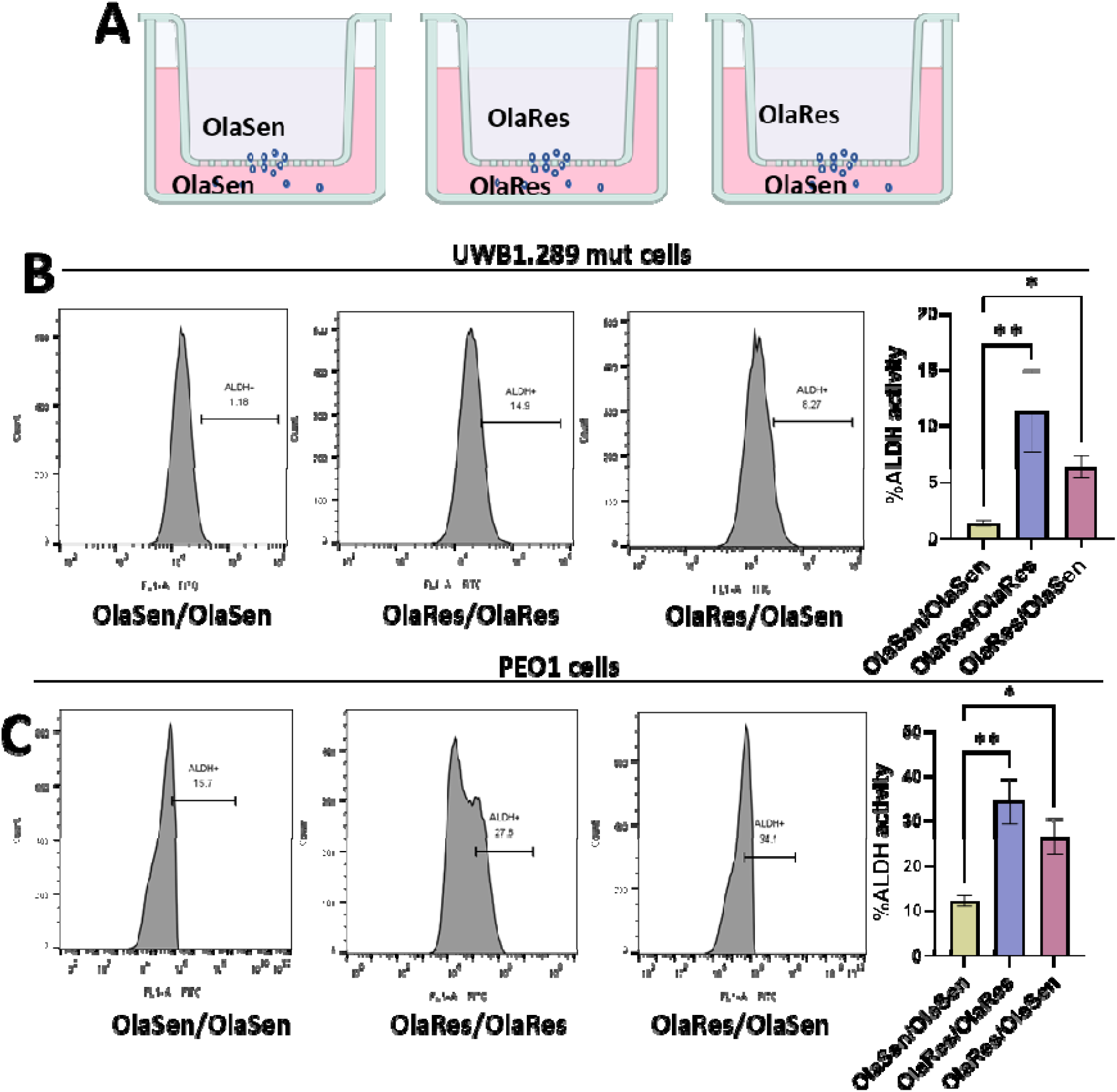
Impact of cell-cell communication on ALDH activity in a co-culture system. A) Schematic representation and quantification of ALDH activity in a transwell co-culture system. Experimental setup: OlaRes cells (high ALDH activity) were cultured in the upper chamber, while OlaSen cells (low ALDH activity) were placed in the lower chamber. Control conditions included OlaRes-OlaRes and OlaSen-OlaSen co-cultures. After allowing cell communication, ALDH activity was measured in (B) UWB1.289 mut and (C) PEO1 cells using flow cytometry. Significance was calculated using one-way ANOVA, with * indicating *P* < 0.05 and ** indicating *P* < 0.01. **Figure 1A** was created with BioRender.com software.

### 3.3. Determining whether EVs from olaparib-resistant cells enhance ALDH activity, sphere-forming ability, olaparib sensitivity, and cell viability in the parental drug-sensitive cells

To investigate whether EVs from olaparib-resistant cells influence cancer stem markers and properties in olaparib-sensitive cells, we assessed ALDH activity, sphere formation, metabolic activity, and apoptosis following EV supplementation from either olaparib-sensitive or -resistant cells. EVs were isolated from both olaparib-sensitive and -resistant UWB1.289 and PEO1 cells, and characterization was performed. Briefly, EVs were isolated from conditioned cell culture medium using a sequential centrifugation, filtration, and ultracentrifugation protocol to ensure the removal of cellular debris and contaminants while preserving vesicle integrity. EVs were then characterized using flow cytometry with a bead-based capture assay and Western blot analysis for purity assessment (see supplemental methods for specific details).

Exposure of UWB1.289 olaparib-sensitive cells with EVs isolated from resistant cells (UWB-OlaRes) increased ALDH activity compared to cells receiving EVs isolated from olaparib-sensitive cells (UWB-OlaSen, Control EVs) (**Figure 2A** **and Figure S9A**). Next, we tested the functional impact of EVs derived from olaparib-resistant cells on sphere-forming capacity. EVs from resistant cells increased the sphere-forming capacity of UWB-OlaSen cells, while spheroid size remained unchanged (**Figure 2B-D**). A similar pattern was observed in PEO1 olaparib-sensitive cells (PEO1-OlaSen), when supplemented with EVs from olaparib-resistant cells (PEO1-OlaRes) (**Figure 2E-H** **and Figure S9B**). We next examined whether EVs from resistant cells influenced the sensitivity of parental cells to olaparib (**Figure 2I**). Exposure of UWB-OlaSen and PEO1-OlaSen cells to EVs from resistant cells UWB-OlaRes and PEO1-OlaRes exhibited higher metabolic activity (**Figure 2J, K**) and decreased levels of apoptotic cell death following olaparib treatment when compared to UWB-OlaSen and PEO1-OlaSen cells exposed to EVs derived from their corresponding olaparib-sensitive cells, as demonstrated by Annexin V staining (**Figure 2L, M and Figure S10A, B**). Interestingly, a similar protective effect was observed in parental drug-sensitive cells supplemented with EVs derived from acutely treated UWB1.289 mut and PEO1 cells, suggesting that even transient exposure to olaparib induces the release of EVs carrying resistance-promoting factors (**Figure S11A**). EVs from olaparib-stressed cells protected against olaparib-induced cell death as determined by metabolic activity (**Figure S11B, C**) and reduced apoptotic cell death (**Figure S11D, E**) in both UWB1.289 mut and PEO1 cells when compared to their control counterparts. Collectively, these results suggest that EVs from olaparib-resistant cells can share stem-like properties, including increased ALDH activity, sphere-forming capacity, and drug resistance.

**Figure 2.**
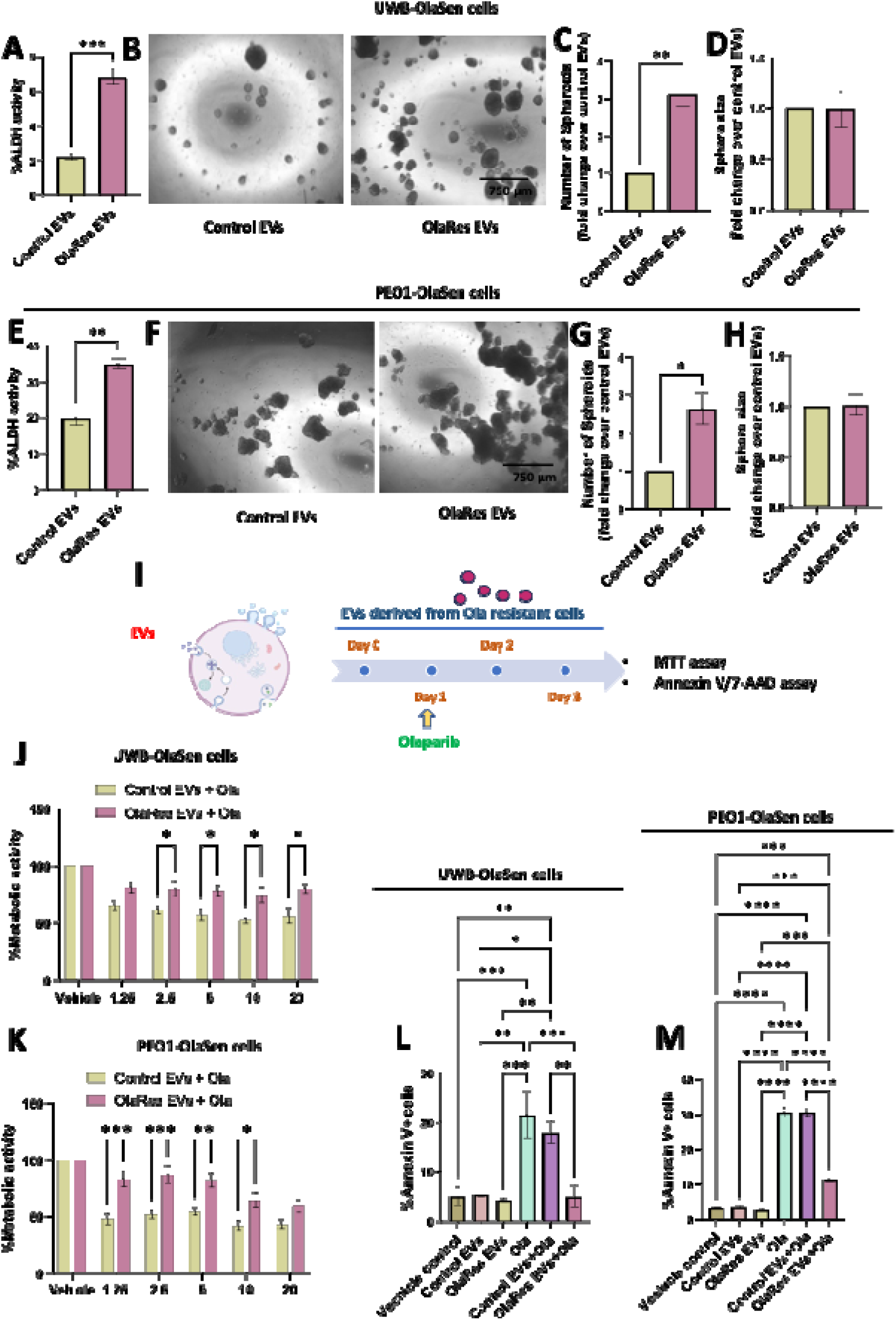
Impact of EVs from olaparib-resistant cells on ALDH activity, sphere-forming ability, olaparib sensitivity, and apoptotic response in drug-sensitive cells. The effect of EVs derived from olaparib-resistant cells on ALDH activity and sphere-forming ability (number of spheroids and sphere size) was examined in (A-D) UWB1.289 mutant olaparib-resistant (UWB-OlaRes) and (E-H) PEO1 olaparib-resistant (PEO1-OlaRes) cells, compared to EVs from control cells (derived from the corresponding parental olaparib-sensitive cell lines). (I) EVs were isolated from UWB-OlaRes and PEO1-OlaRes cells. UWB-OlaSen and PEO1-OlaSen parental cells were treated with EVs at a concentration of 2.5 µg/mL or vehicle and cultured for 24 hours (Day 0), before treatment with olaparib, with daily EV supplementation for 3 days. Metabolic activity of (J) UWB1.289 mut and (K) PEO1 parental cells was assessed 72 hours post-treatment using an MTT assay. Similarly, apoptotic cell death in (L) UWB1.289 mut and (M) PEO1 parental cells was evaluated using annexin V binding assays. Significance was calculated using *t*-test, one-way, and two-way ANOVA, with * indicating *P* < 0.05, ** indicating *P* < 0.01, *** indicating *P* < 0.001, and **** indicating *P* < 0.0001. **Figure 2I** was created with BioRender.com software.

Independently, we examined whether EVs from drug-treated A2780 cells or ALDH+ spheroid-enriched A2780 cells, both enriched for CSCs, could similarly impact metabolic activity in recipient parental cells **(Figure S12A)**. A2780 monolayer cells were pre-treated with 5 µM of carboplatin (**Figure S12B**) or olaparib (**Figure S12C**), and then EVs were collected and isolated after 72 hours. Recipient cells were primed with EVs at a concentration of 2.5 µg/ml 24 hours (Day 0) before treatment with carboplatin or olaparib, with daily EV supplementation (2.5 µg/ml) for 3 additional days. Metabolic activity was assessed 72 hours post-treatment using an MTT assay. Exposure to these EVs inhibited the reduction in metabolic activity induced by subsequent drug exposure compared to their vehicle controls. Similarly, EVs derived from ALDH activity-positive spheroid-enriched A2780 cells (**Figure S12D**) inhibited the reduction of metabolic activity in recipient cells induced by drug treatment. Together, these data reinforced the hypothesis that EVs derived from stem-like populations transmit survival signals to neighboring cells.

### 3.4. Assessing whether EVs isolated from patient-derived organoids (PDOs) established from patients with recurrent OvCa could enhance stem-like properties

Organoid culture conditions are reported to be enriched for cells with stem-like properties (Condello et al., 2015). Thus, we postulated that EVs from PDOs generated from patients diagnosed with recurrent HGSOC could promote stem-like properties in drug-sensitive cells. To test this, two PDO models were utilized. Both 17-121 and VCRB 330 PDOs were derived from patients with recurrent OvCa after having received one or more lines of therapy, implying they developed some level of resistance to prior treatments. Exposure to EVs from 17-121 and VCRB330 PDO conditioned media using the same strategy and concentrations described above (section 3.3) increased the percent of cells with ALDH activity in UWB1.289 mut cells (**Figure 3A and Figure S13A**), suggesting a role of EVs in enhancing a cancer stem-like marker. This increase in ALDH activity after exposure to EVs from the PDO was concurrent with an increase in sphere-forming capacity (**Figure 3B-D**). Next, we examined whether PDO-derived EVs affected the sensitivity of UWB1.289 cells to olaparib treatment. A schematic overview of the experiment is shown in **Figure 3E**. Exposure of EVs from the PDOs offset some of the loss in metabolic activity in olaparib-treated control cells, indicating a protective effect against drug-induced cytotoxicity (**Figure 3F**). This survival advantage was further confirmed by annexin V assays, which showed a significant reduction in olaparib-induced apoptotic cell death in PDO EV-exposed cells (**Figure 3G and Figure S13B**). These findings suggested that EVs from conditioned media from the OvCa PDOs contribute to increased stem-like properties.

**Figure 3:**
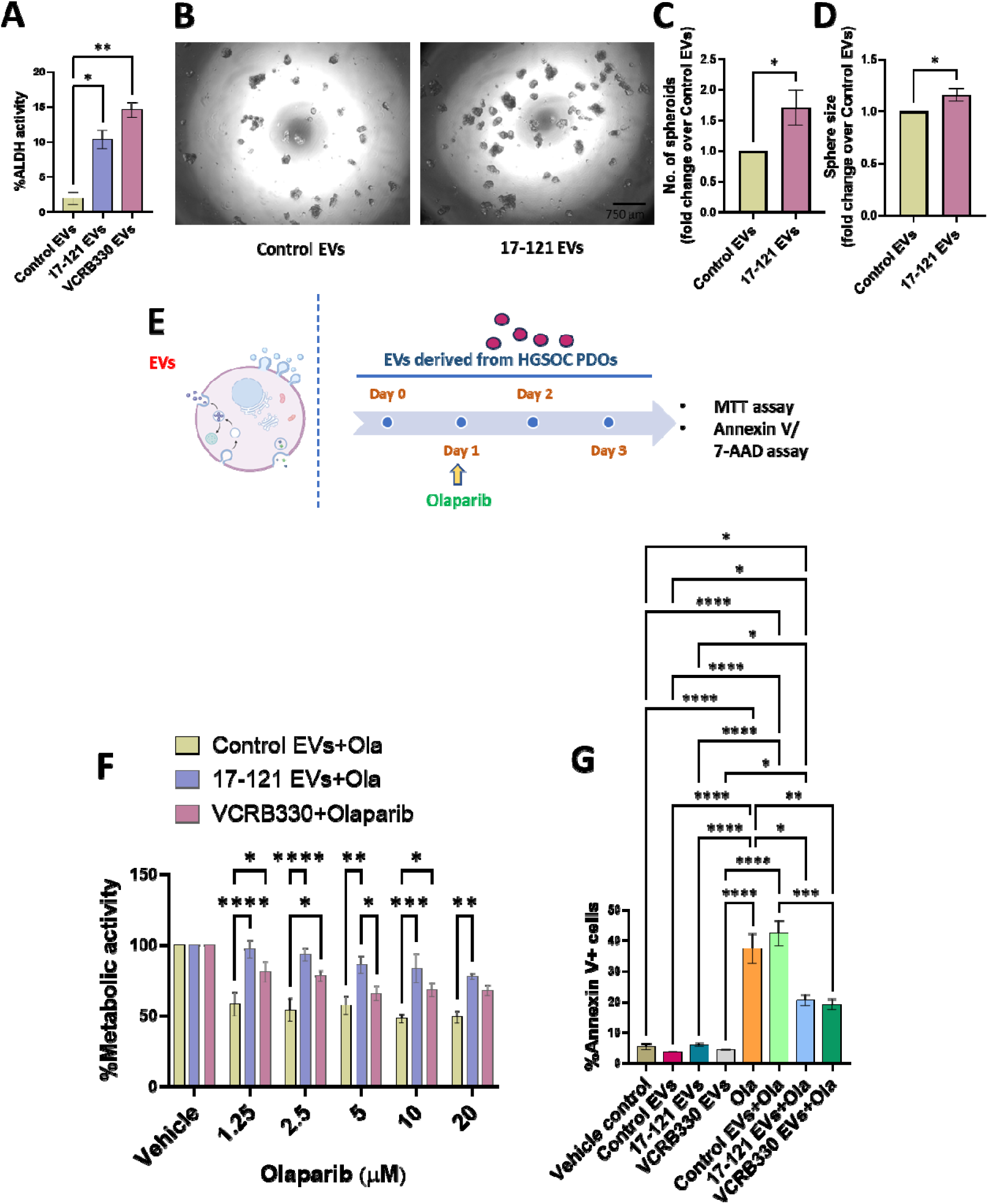
EVs from HGSOC patient-derived organoids (PDOs) influence stemness, olaparib sensitivity, and viability in UWB1.289 mut cells. EVs were isolated from HGSOC PDOs 17-121 and VCRB330. (A) UWB1.289 mutant (mut) cells were treated with EVs from PDOs 17-121 and VCRB330 at a concentration of 2.5 µg/ml for 72 hours, followed by assessment of ALDH activity using flow cytometry. (B) Sphere-forming ability was assessed to check the impact of EVs from 17-121 PDO in UWB1.289 mut cells, with the (C) number of spheroids and (D) sphere size measured using ImageJ analysis. (E) UWB1.289 mut cells were treated with EVs (2.5 µg/ml) or vehicle for 24 hours (Day 0), followed by additional EV supplementation on Day 1 and for two more days. Olaparib treatment was administered on Day 1 only. (F) Metabolic activity was measured 72 hours post-treatment using an MTT assay. (G) Apoptotic cell death was assessed by annexin V binding assays. Statistical significance was determined using *t*-test, one-way, and two-way ANOVA, with *P < 0.05, **P < 0.01, ***P < 0.001, and ****P < 0.0001. **Figure 3E** was created with BioRender.com software.

### 3.5. Assessing whether EVs from olaparib-resistant cells can minimize evidence of DNA damage in olaparib-sensitive BRCA-mutant cells

We previously demonstrated that PARPi-enriched ovarian CSCs display enhanced DNA repair properties (Bellio et al., 2019a). To assess whether EVs from OlaRes cells influenced the number of cells displaying DNA damage, we performed a Comet assay. The Comet assay revealed that exposure to EVs from resistant cells minimized the number of cells displaying DNA damage, as evidenced by a marked decrease in the number of tails, denoting damaged DNA observed in both UWB1.289 (**Figure 4A, B**) and PEO1 (**Figure 4C, D**) cells. Together, these findings suggested that EVs from olaparib-resistant cells transfer protective signals that either prevent DNA damage or enhance DNA repair, mitigating the toxic effects of olaparib.

**Figure 4:**
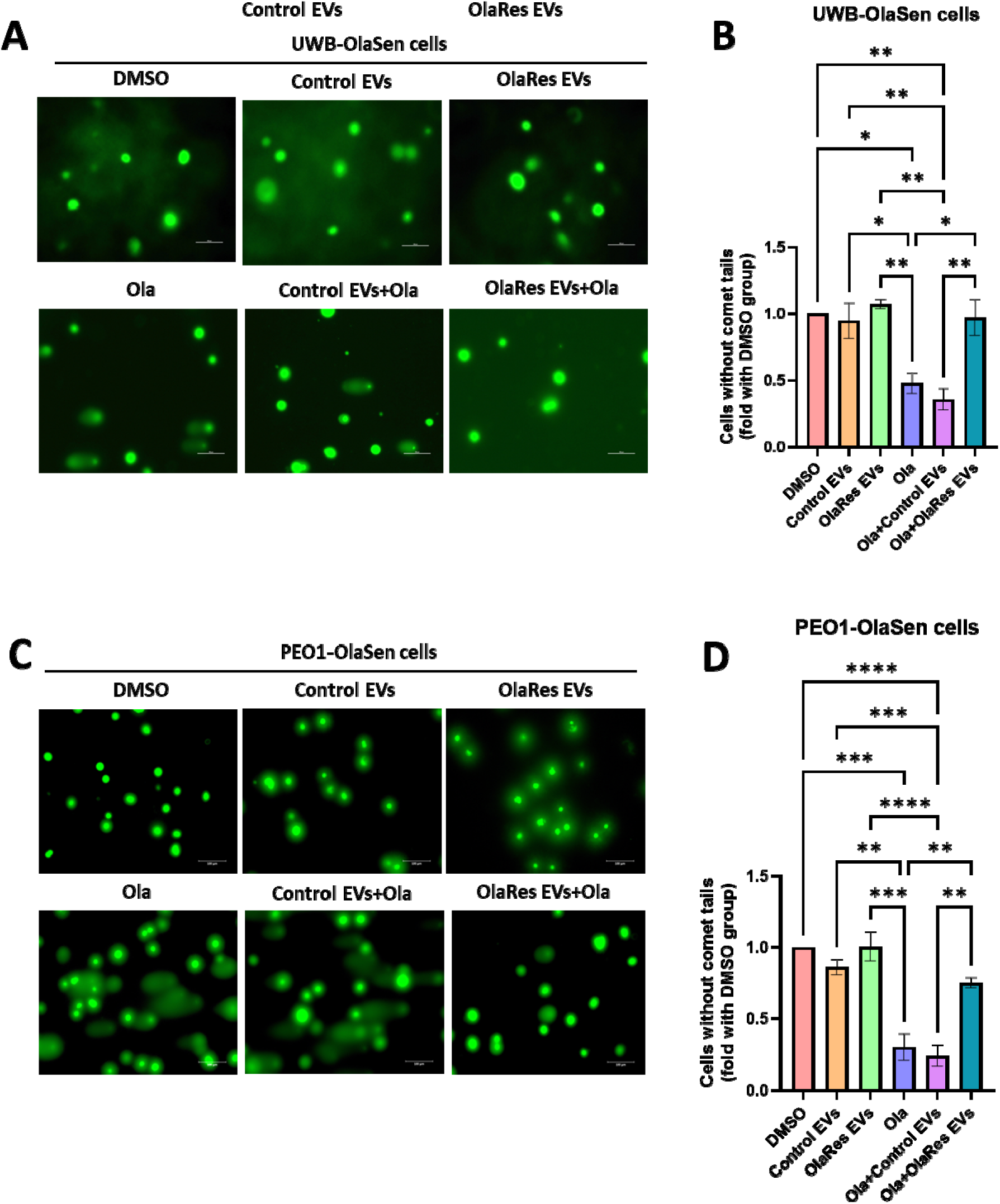
Effect of EVs on DNA damage in UWB1.289 mut and PEO1 cells. Both (A, B) UWB1.289 mut and (C, D) PEO1 cells were treated with EVs (2.5 ug/ml) from PEO1 OlaRes or control cells, followed by 10 μM olaparib (Ola) or DMSO. EVs (2.5 ug/ml) were replenished daily for 72 hours. DNA damage was assessed using a comet assay, and images were captured at 200× magnification. The cells without comet tails were calculated and quantified. Statistical significance was determined using one-way ANOVA, with *P < 0.05, **P < 0.01, ***P < 0.001, and ****P < 0.0001.

### 3.6. EVs from OlaRes cells promote EZH2-mediated epigenetic reprogramming and DNA damage response signaling in OvCa cells

EZH2, which is often linked to stemness, is elevated in OvCa (Cardenas et al., 2016; Jones et al., 2018; Reid et al., 2021; Rizzo et al., 2011). Furthermore, there is evidence of a mechanistic link between PARP and EZH2 (Zhang et al., 2022). Therefore, our next objective was to assess the impact EVs derived from OlaRes cells have on EZH2 expression and potential associated epigenetic modifications in OlaSen counterparts. We showed that EVs from UWB-OlaRes cells increased EZH2 expression in UWB-OlaSen cells (**Figure 5A**) compared to the control EVs derived from the olaparib-sensitive cells. A similar effect was observed in PEO1-OlaSen cells when treated with EVs from PEO1-OlaRes cells (**Figure 5B and Figure S14A**) compared to their control. The upregulation of EZH2 was concordant with an increase H3K27me3 levels in both UWB1.289 mut (**Figure 5C**) and PEO1 (**Figure 5D and Figure S14B**) olaparib sensitive cell lines compared to their respective controls, suggesting the EVs from the resistant cells had a potential role in epigenetic reprogramming of the drug-sensitive cells.

**Figure 5.**
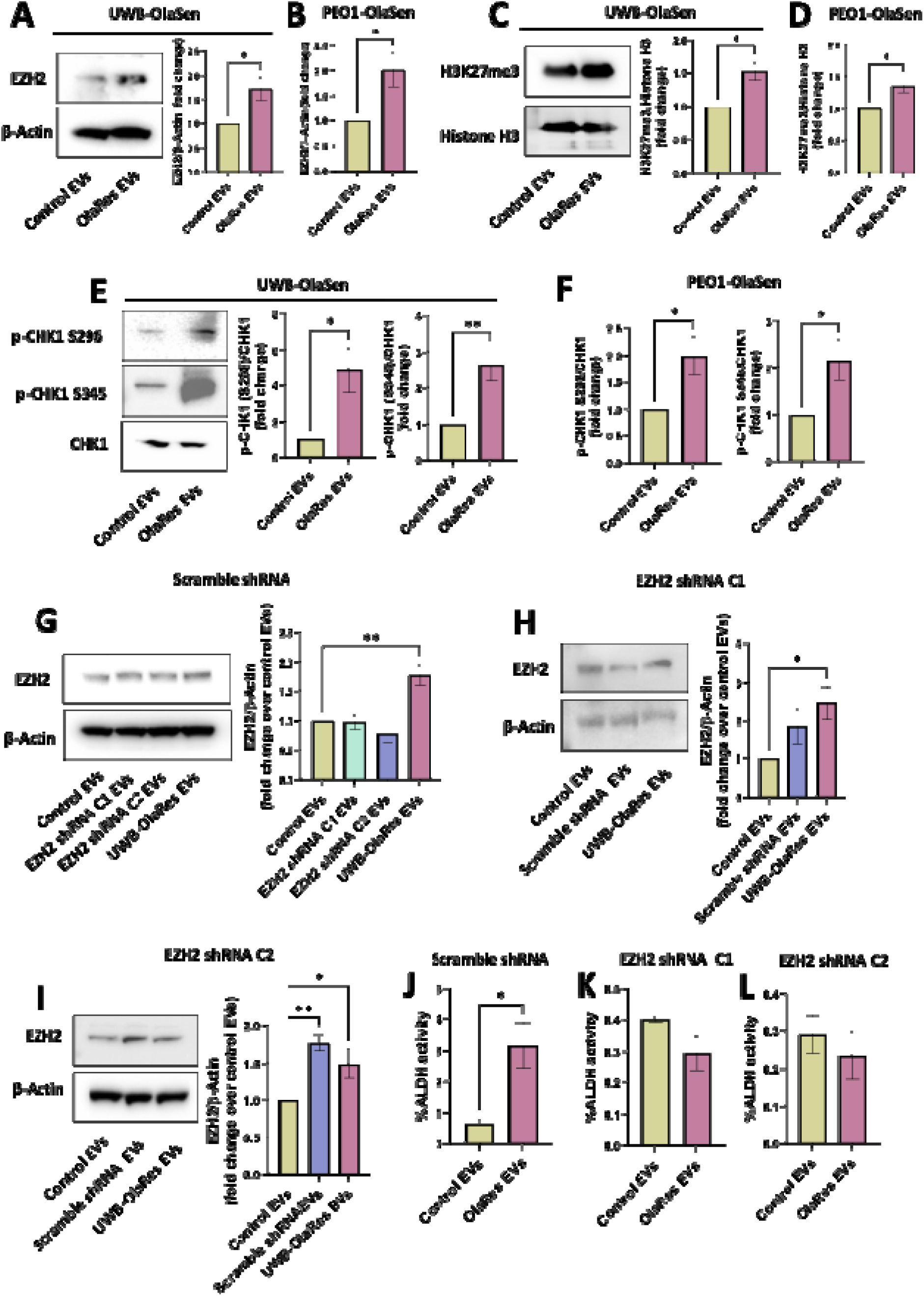
Impact of EVs from olaparib-resistant cells on EZH2 expression, H3K27me3 levels, CHK1 phosphorylation, and stemness in olaparib-sensitive and EZH2 knockdown cells. Olaparib-sensitive (UWB-OlaSen and PEO1-OlaSen) cells were treated with EVs derived from olaparib-resistant UWB1.289 (UWB-OlaRes) and PEO1 (PEO1-OlaRes) cells or control cells. After 72 hours of incubation, western blot analysis was performed to assess (A, B) EZH2 and (C, D) H3K27me3 expression in UWB-OlaSen and PEO1-OlaSen cells, respectively, following EV treatment. CHK1 phosphorylation at S296 and S345 was examined by western blotting after 30 minutes of EV incubation in (E) UWB-OlaSen and (F) PEO1-OlaSen cells, respectively. The effect of EVs from UWB-OlaRes or control cells on ALDH activity was evaluated using flow cytometry in (G) Scramble shRNA, (H) EZH2 shRNA C1, and (I) EZH2 shRNA C2 cells. (J) Western blot analysis of EZH2 expression in scramble shRNA cells treated with EVs from EZH2 shRNA C1, C2, or UWB-OlaRes cells. Western blot analysis of EZH2 expression in (K, L) EZH2 shRNA C1 and C2 cells following treatment with EVs from scramble shRNA, UWB-OlaRes, or control cells. Statistical significance was determined using *t*-test and one-way ANOVA, with *P < 0.05, **P < 0.01, and ***P < 0.001 (n ≥ 3 independent biological sets).

Independent of EZH2 canonical histone modifications, it was previously reported that EZH2 has the potential to bind and mediate transcription and activation of CHK1 signaling in OvCa cells, promoting drug resistance and maintaining stem-like features (Wen et al., 2021). We investigated the effect of EVs from OlaSen and OlaRes on CHK1 phosphorylation, a marker of CHK1 activation in olaparib-sensitive cell lines. We observed that exposure to EVs from OlaRes cells increased CHK1 phosphorylation at S296 and S345 in UWB1.289 mut (**Figure 5E**) and PEO1 (**Figure 5F and Figure S14C**) cells, again revealing that EVs from OlaRes cells could also promote signaling via non-canonical DNA damage response pathways. This aligned with Wen et al., who reported that EZH2 bound and initiated gene transcription of CHK1 and promoted CHK1 phosphorylation (Wen et al., 2021). EVs derived from 17-121 and VCRB330 PDOs unchanged the expression of EZH2 but enhanced its associated histone modification, H3K27me3, in UWB1.289 mut cells compared to EVs from drug-sensitive UWB1.289 mut cells (**Figure S15A–D**), suggesting that EVs from PDOs derived from recurrent tumors may contribute to epigenetic alterations that promote a more resistant phenotype. Additionally, EVs from recurrent PDOs selectively increased CHK1 phosphorylation at S296 but not at S345 (**Figure S15E, F**), indicating a potential role of EV-mediated signaling in regulating post-translational modifications that could influence therapeutic response and cellular adaptability.

We next assessed whether EZH2 might be present in the EVs recovered from the drug-resistant cells, implicating a direct role. Using western blotting techniques, we found no evidence of EZH2 protein in the EVs from any of the cell lines we analyzed. Others have shown that *EZH2* is present in EVs, and that cells exposed to these EVs exhibit an increase in *EZH2* in recipient cells, implying that the mRNA is transferred (Villasante et al., 2021; Villasante et al., 2016).

While we showed *EZH2* was present in EVs derived from both olaparib-sensitive and resistant UWB1.289 cells using RT-qPCR, we observed no differences in the levels of *EZH2* in the EVs from the sensitive and resistant lines. In contrast to the findings of others (Villasante et al., 2021; Villasante et al., 2016), we did not observe any consistent differences in EZH2 levels in cells exposed to EVs from sensitive and resistant cells.

We next tested the impact of endogenous levels of EZH2 in the EV donor cells and what impact it might have on the capacity of olaparib-sensitive cells to acquire more stem-like properties initiated by EV exposure. Using our scrambled and shRNA EZH2 KD lines, we demonstrated that only exposure to EVs from UWB-OlaRes cells increased EZH2 levels in scrambled shRNA cells, an effect that was not observed when EVs originated from EZH2 shRNA C1 and C2 cells (**Figure 5G**), suggesting that some component of the cargo carried by EVs from the resistant cells contributes to the upregulation of EZH2 in the recipient cells. However, despite EZH2 knockdown, EVs from UWB-OlaRes cells still enhanced EZH2 expression in EZH2 shRNA C1 cells (**Figure 5H**), indicating the presence of a factor independent of EZH2 may be actively playing a role. Additionally, EVs from both scramble shRNA and UWB-OlaRes cells increased EZH2 expression in EZH2 shRNA C2 cells compared to their control (**Figure 5I**). Collectively, these findings suggest that the transfer of an EZH2-associated regulatory factor may also contribute to the plasticity, DNA damage response, and stemness-like features. In support of these findings, we showed that exposure of the sensitive cells with EVs from OlaRes cells significantly increased ALDH activity in UWB1.289 mutant cells hosting a scrambled shRNA (**Figure 5J and Figure S16A**). However, this effect was not seen in cells with reduced levels of EZH2 in the EZH2 KD lines (**Figure 5K, L, and Figure S16B, C**), suggesting the need for a sufficient threshold level of EZH2 to observe an increase in ALDH activity. These findings further imply that it is not just EZH2 mediating the transfer of stem-like properties.

### 3.7. Exploring the potential for HOTAIR-mediated EV transfer and its role in transferring stemness and drug resistance

The mechanism(s) through which EVs regulate stemness, drug resistance, and DNA damage repair in OvCa remain unclear. HOTAIR, a long noncoding RNA (lncRNA), was identified as a regulator of OvCa stemness and drug resistance by activating EZH2 and contributing to PRC2-mediated epigenetic modifications (Özeş et al., 2017). To explore its role in olaparib resistance, we examined baseline HOTAIR expression in both OlaSen and OlaRes OvCa cells. OlaRes cells had higher HOTAIR expression in both UWB1.289 mut and PEO1 cell lines (**Figure 6A**) relative to the OlaSen UWB1.289 and PEO1 cells. Additionally, using RT-PCR, we showed that PEO1-OlaRes EVs contained significantly higher HOTAIR levels than their OlaSen counterparts, whereas there was no difference in HOTAIR expression in UWB1.289 EVs (**Figure 6B**). These results confirmed that OlaRes-derived EVs transport HOTAIR as part of their cargo, suggesting the potential for horizontal transfer of this lncRNA between CSC/drug-resistant cells and non-CSCs. HOTAIR expression increased in the OlaSen cells following OlaRes EV exposure (**Figure 6C**) compared to their appropriate controls. These findings suggest that EV-mediated HOTAIR transfer may mediate EZH2’s effect in enhancing stem-like properties and contributing to drug resistance in OvCa cells.

**Figure 6.**
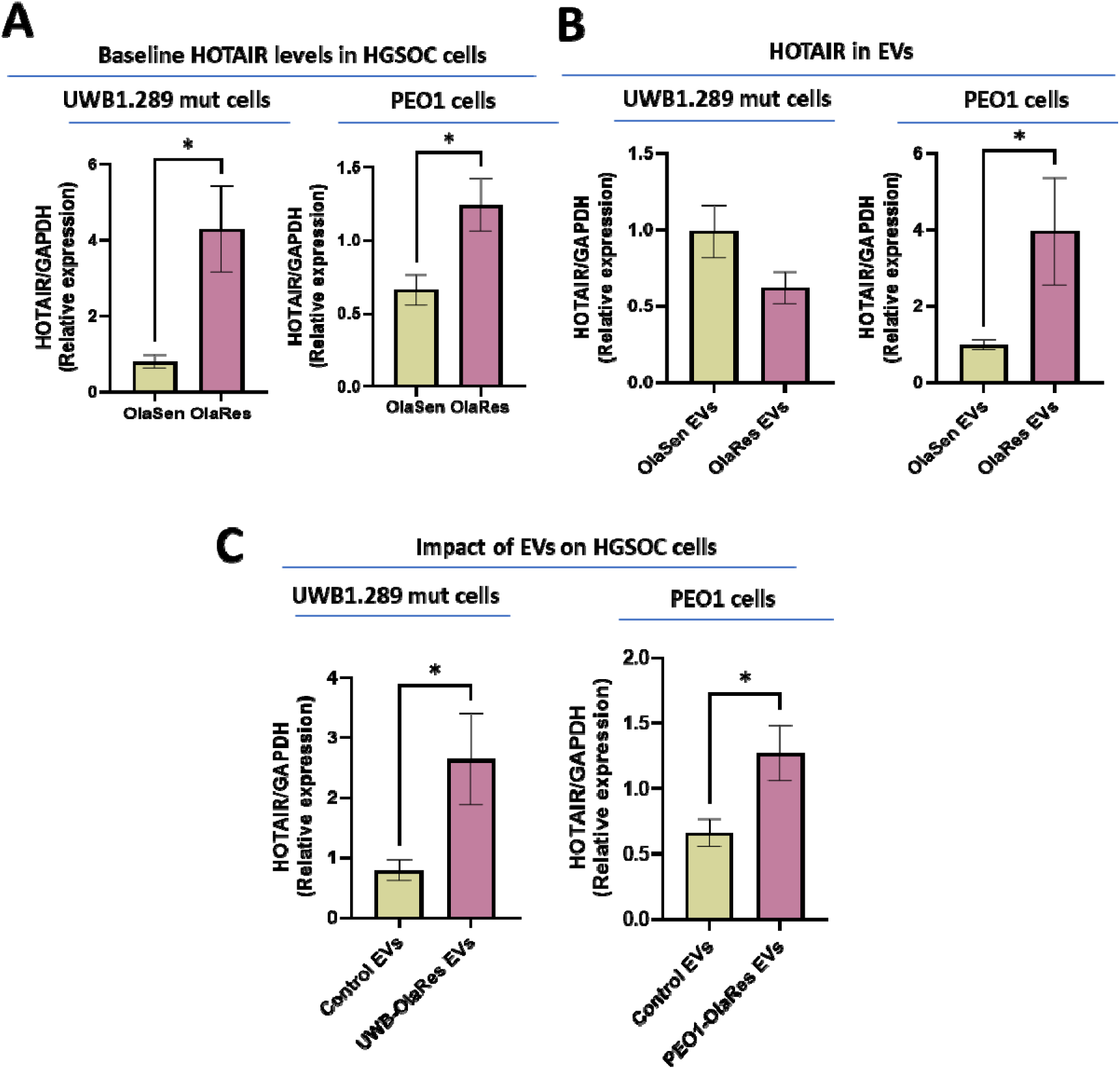
Quantification of HOTAIR expression in OlaSen and OlaRes cell lines and EVs. (A) Quantification of HOTAIR levels in OlaSen and OlaRes UWB1.289 mutant (mut) and PEO1 cells, determined by RT-PCR. (B) HOTAIR expression levels in EVs isolated from OlaSen and OlaRes cells of UWB1.289 mut and PEO1, assessed via RT-PCR. (C) OlaSen UWB1.289 mut and PEO1 cells were supplemented with EVs derived from OlaRes cells and incubated for 6 hours. HOTAIR levels in recipient cells were measured by RT-PCR post-incubation. Statistical significance was determined using a *t*-test, with *P < 0.05.

### 3.8. Impact of EVs from OlaRes cells on apoptotic priming and mitochondrial network remodeling in UWB1.289 mut cells

Drug resistance in OvCa is often mediated at the mitochondrial level by a balance of pro- and anti-apoptotic proteins (Al-Alem et al., 2019a). Given that we demonstrated ovarian CSC and drug-resistant cells can resist olaparib-induced cell death, we investigated whether EVs derived from OlaRes cells had an impact on apoptotic priming, which is defined as the proximity of cells to the apoptotic threshold. Changes in apoptotic priming were previously shown to alter the sensitivity of OvCa cells to cancer therapeutics *in vitro* and in the clinical setting (Ni Chonghaile et al., 2011; Stover et al., 2019). Apoptotic priming was measured using BH3 profiling (Fraser et al., 2019), which detects cytochrome c release as a readout for initiation of apoptosis in response to increasing concentrations of pro-apoptotic BIM and BID BH3 peptides (Fraser et al., 2019).

Baseline analysis revealed that UWB1.289 cells were responsive to moderate (10 μM) and high (100 μM) doses of these pro-apoptotic peptides, but not to low (1 μM) doses, as indicated by cytochrome c release upon exposure to these treatments (**Figure 7A**). These results demonstrated that these cells were moderately primed for apoptosis. BH3 profiling also measures cellular dependence on anti-apoptotic proteins by measuring apoptosis initiation in response to BH3 peptides that bind and inhibit specific anti-apoptotic proteins (Fraser et al., 2019). BH3 profiling analysis revealed that UWB1.289 cells exhibited dependence on anti-apoptotic proteins BCL-2, BCL-W, and/or BCL-XL, as indicated by their sensitivity to the BAD peptide, which selectively inhibits these proteins (Fraser et al., 2019). The dependence on BCL-XL was further classified as mild due to the low response observed with the HRK peptide, which specifically targets BCL-XL. No significant dependence on MCL-1 was detected.

**Figure 7.**
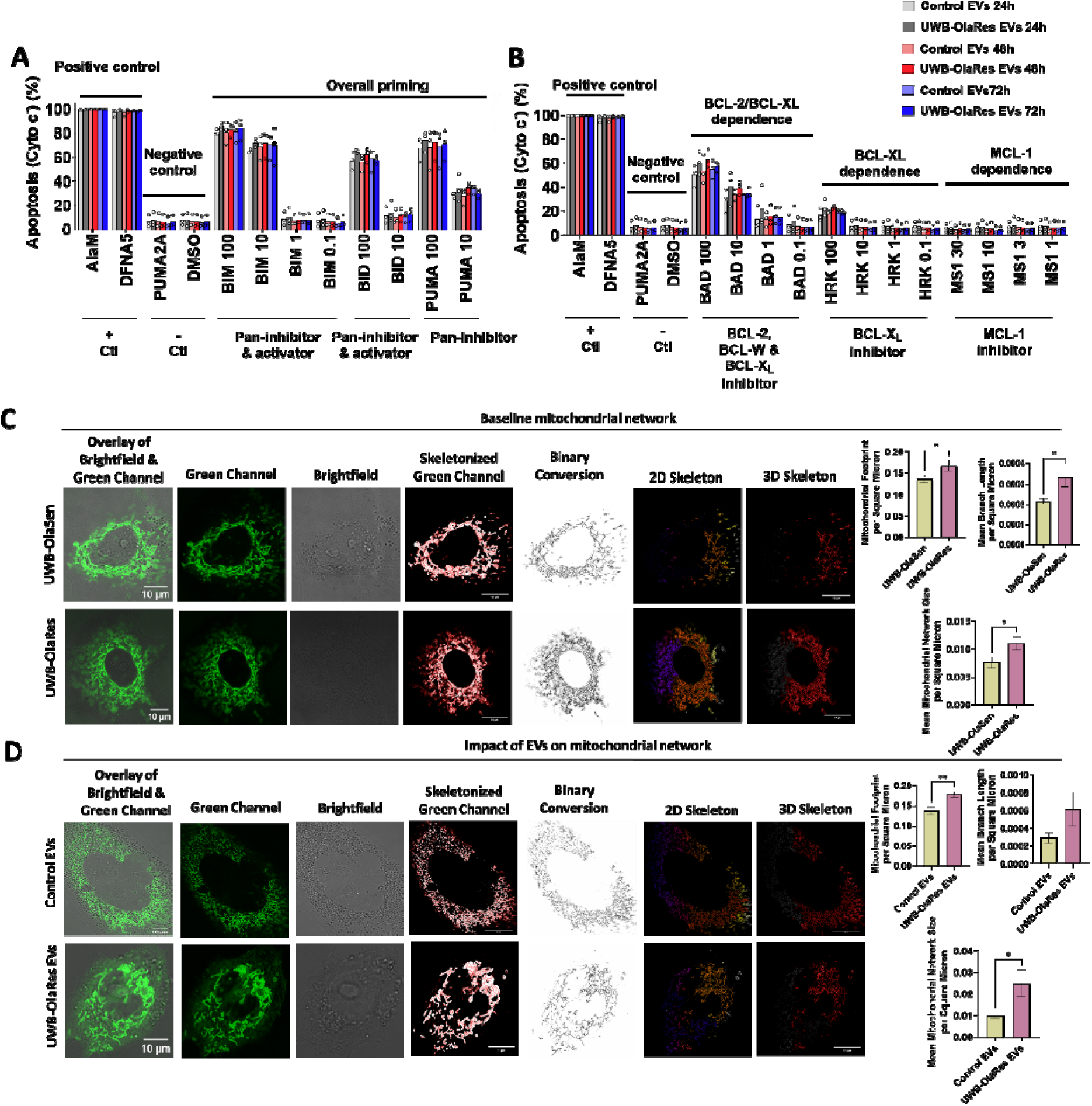
Impact of OlaRes-derived EVs on apoptotic priming and mitochondrial network remodeling in OvCa cells. UWB1.289 mut OlaSen cells were treated with EVs derived from OlaRes cells for 24, 48, and 72 hours, followed by BH3 profiling using flow cytometric analysi to assess apoptotic (A) priming and (B) dependencies. Mitochondrial network analysis wa performed by staining cells with MitoTracker Green, capturing images using the 63× objective of a Zeiss LSM 880 Confocal Fluorescence Microscope, and quantifying mitochondrial parameters, including mitochondrial footprint, mean branch length, and mean mitochondrial network size, using ImageJ. Image panels include the overlay of the fluorescent green channel and the brightfield with a 10 µm scale bar, the fluorescent green channel, the brightfield, the skeletonized image, the binary conversion, the 2D network, and the 3D network. (C) Baseline mitochondrial morphology and network characteristics in UWB-OlaSen and OlaRes cells. (D) Mitochondrial network analysis in OlaSen cells following supplementation with OlaRes-derived EVs, showing changes in mitochondrial morphology and dynamics. Statistical significance was determined using a *t*-test and two-way ANOVA, with *P < 0.05 and **P < 0.01.

Following EV exposure, no significant difference in apoptotic priming or dependencies on pro-survival BCL-2 family was observed in UWB1.289 cells treated with OlaRes EVs compared to those exposed to control EVs. Notably, cells treated with OlaRes EVs exhibited enhanced dependence on BCL-2 and BCL-XL, suggesting that these EVs modulate apoptotic signaling pathways. These effects were observed across multiple time points, with moderate but reproducible differences in apoptotic priming and BCL-2, BCL-W, and BCL-XL dependency (**Figure 7B**). These findings suggest that OlaRes EVs may alter the sensitivity of recipient cells to undergo apoptosis, potentially influencing cell survival mechanisms in the context of ovarian cancer therapy resistance. Moreover, the findings suggested that ovarian CSCs or non-CSCs taking on stem-like features via EVs may be more vulnerable to treatments targeting specific BCL-2 family members (Fraser et al., 2019; Sarosiek and Wood, 2023).

Mitochondrial network dynamics have been implicated in both stemness and therapy resistance (Fan et al., 2019; Ni Chonghaile et al., 2011; Tan et al., 2025). Thus, we evaluated whether EVs affected the mitochondrial network. The analysis revealed distinct alterations in mitochondrial morphology. At baseline, OlaRes UWB1.289 mut cells exhibited an increased mitochondrial footprint, mean branch length, and mitochondrial network size compared to OlaSen cells (**Figure 7C**). To further validate these findings, proMF2 nanoparticles (NPs) were used as a positive control and compared to untreated UWB1.289 mut cells (**Figure S17**). Importantly, EVs derived from UWB-OlaRes cells enhanced mitochondrial footprint, mean branch length, and mitochondrial network size in recipient OlaSen cells compared to control EVs from UWB-OlaSen cells (**Figure 7D**). These findings indicate that OlaRes EVs contribute to mitochondrial remodeling, which may influence cellular metabolism and apoptotic susceptibility in OvCa cells.

## 4. Discussion

Recurrent OvCa remains an important clinical challenge. PARP inhibitor-based strategies have had some success, yet many patients develop resistance. While others and we provided evidence that OvCa CSCs are enriched post-treatment with cytotoxics or more targeted strategies like PARPi (Al-Alem et al., 2019b; Bellio et al., 2019a; Nacarelli et al., 2020; Wang et al., 2014; Wang et al., 2018; Zong and Nephew, 2019), there is mounting evidence that drug-sensitive cells can demonstrate some level of plasticity and/or take on more CSC-like properties (Bapat et al., 2010; Berry and Bapat, 2008; Kusumbe et al., 2009). Our objective was to provide evidence that this plasticity or dedifferentiation is due in part to communication between CSCs or drug-resistant tumor cells with drug-sensitive tumor cells. Moreover, we hypothesized that this information exchange was mediated, in part, by EVs and their cargo, driving some epigenetic modifications. Our results provide evidence that stem-like properties can be shared among tumor cells and that EZH2 can partially mediate these properties. In this study, we confirmed previous findings that treatment of OvCa cells with a PARPi results in an enrichment of CSCs (Bellio et al., 2019a; Bellio et al., 2019b). This enrichment was negated with the EZH2 inhibitor, GSK126. Subsequent analysis showed that exposure to PARPi resulted in an increase in EZH2, not unlike what others showed following treatment with a cytotoxic (Wang et al., 2021). These findings align with those of others who found that targeting other pathways linked to the promotion or maintenance of CSCs will negatively impact OvCa cell survival and tumor growth. These include, but are not limited to, Notch or Hedgehog or other factors like OCT4, WNT, SOX2 or NANOG (Cioffi et al., 2015; Groeneweg et al., 2014a; Groeneweg et al., 2014b; Huang et al., 2015; McCann et al., 2011; Peng et al., 2010; Robinson et al., 2021; Zong et al., 2020).

We initially focused on EZH2 as it was shown to be elevated in OvCa (Cardenas et al., 2016; Jones et al., 2018; Reid et al., 2021; Rizzo et al., 2011). One report suggested EZH2 expression could be a predictor of platinum resistance (Reid et al., 2021). Moreover, there was mounting evidence of a mechanistic link between PARP and EZH2 (reviewed by Zhang et al.) (Zhang et al., 2022) and several preclinical studies where they combined PARPi with EZH2 inhibitors or used a knockdown strategy for reduction of EZH2 along with PARPi in OvCa *in vitro* and *in vivo* models (Karakashev and Zhang, 2020; Sun et al., 2021). We were particularly focused on how CSCs might escape these types of strategies and whether the expected resistance would be due in part to some level of protectiveness initiated by the CSCs and/or drug-resistant populations. Our simple co-culture experiments highlighted how one tumor cell line, with a greater percentage of cells exhibiting ALDH activity, indicating a higher percentage of CSCs, could influence another tumor cell line with less activity, thereby increasing its ALDH activity above its baseline. The lack of direct cell-to-cell contact and the size of the membrane separating the two cell types reinforced our hypothesis that the signals being communicated might be exchanged via EVs, at least in part. This led us to develop multiple models to test the contribution of EVs from CSCs or drug-resistant cells to drug-sensitive cells. To this end, we isolated and characterized EVs from drug-resistant and sensitive cells and assessed their impact on the drug-sensitive cells, including whether they differentially promoted CSC markers and/or phenotypes and how they may alter response to treatment.

As mentioned, the plasticity of cells refers to the ability of a specific population of cells to change between different phenotypical states. More specifically, it is proposed that the more differentiated tumor cells have the potential to shift to a CSC phenotype or vice versa under an appropriate stimulus. It is anticipated that genetic, epigenetic, pharmacologic and/or microenvironmental changes can trigger the bidirectional interconversion between stem and non-stem-like states (van Neerven et al., 2016). The integration of CSCs, clonal evolution and potential for bidirectional interconversion provides an alternative survival mechanism for the tumor. We hypothesize that recurrent treatment resistant OvCa is a byproduct of all these models contributing to the heterogeneity of the disease, which is further modified by clinical treatment regimens. Moreover, we predict that treatment induced enrichment of CSCs promotes exchange of these properties via EVs.

EVs, including small EVs and microparticles, are implicated in communicating drug-resistance in a number of solid tumors (Namee and O’Driscoll, 2018; Samuel et al., 2017; Xavier et al., 2020), including OvCa (Dai et al., 2024; Samuel et al., 2018; Tian et al., 2022). Most of the focus has been on EVs from stromal cells and adipose cells and their role in promoting drug resistance and/or supporting ovarian CSCs. However, others have shown EVs can carry cargo that can negatively impact drug resistance in OvCa (Suzuki et al., 2023; Wang et al., 2022).

EZH2 has been linked to the long noncoding HOX antisense intergenic RNA (HOTAIR). HOTAIR is reported to promote cell cycle progression and mediate DNA damage repair by interacting with EZH in glioblastoma (Yang et al., 2024). HOTAIR also promotes gefitinib resistance through the modification of EZH2 in small-cell lung cancer (Li et al., 2021). HOTAIR reprograms chromatin state to promote metastasis in preclinical breast cancer models (Gupta et al., 2010). Wang et al. demonstrated that HOTAIR was upregulated in ovarian CSC relative to non-CSC. They found that ectopic expression of HOTAIR enriched the ALDH-positive cell population. Similarly, EZH2 overexpression increased colony and sphere-forming capacity.

Pretreatment of OVCAR3 cells *in vitro* with the peptide nucleic acid -PNA3 reduced tumor initiation, growth, and CSC frequency, further emphasizing their interaction in mediating stem-like properties(Wang et al., 2021). While we did not detect EZH2 protein in the small EVs, like others, we detected *EZH2* in EVs by PCR. However, we did not discern any differences in *EZH2* levels in EVs isolated from conditioned media from olaparib-sensitive and -resistant cells, nor did we see any consistent increase in *EZH2* levels in the recipient cells following EV exposure. This contrasts with other models where an increase in *EZH2* was observed in the recipient cells following EV exposure, albeit in bioengineered or stromal cell lines (Villasante et al., 2021; Villasante et al., 2016). In the absence of EZH2, it was not unrealistic to test whether HOTAIR might be present in EVs, and that the transfer of EVs from drug-resistant cell populations, which are enriched for ALDH-positive cells, would result in an increase in HOTAIR in the recipient cells. In fact, we did observe that the olaparib-resistant UWB1.289 and PEO1 cells both displayed higher levels of HOTAIR compared to their drug-sensitive controls. Exposure of the olaparib-sensitive cells to EVs from resistant cells resulted in an increase in HOTAIR. Given that ectopic expression of HOTAIR promotes stem-like features, it is highly possible that HOTAIR is one of many factors exchanged between CSCs or drug-resistant cells and non-CSC or drug-sensitive cells, thereby promoting stemness. Others documented that lncRNAs are key regulators of epithelial and mesenchymal cell plasticity and stemness (Yuan et al., 2025). Moreover, there are likely more microRNAs that may have a similar effect in promoting drug resistance, plasticity, and stemness in OvCa (Chong, 2024; Dey Bhowmik et al., 2024).

Independently, we demonstrated EVs from OlaRes OvCa cells had the capacity to transform the mitochondrial dynamics of OlaSen OvCa cells. This presumably confers a survival advantage as networked mitochondria are resistant to apoptosis. EV signaling alters cell phenotypes at multiple levels, including drug resistance, stemness, and organelle dynamics. Long noncoding RNAs and miRNAs are known to influence mitochondrial function (Fan et al., 2019; Sun et al., 2022), expanding the scope of study on how they may mediate plasticity in OvCa cells.

Collectively, these data reinforce the need for combination strategies, targeting tumor cells as they present and what they can become under the selection pressure influenced by the microenvironment, different treatment regimens, and the tumor cell’s ability to undergo continued epigenetic changes, further driving tumor cell evolution. Given the high feasibility of serial blood sampling for EV profiling, the novel biological insights generated here may facilitate earlier “go-no go” decision-making and improve (pre) clinical outcomes.

## AUTHOR CONTRIBUTIONS

VP, investigation; methodology; data curation; formal analysis; acquisition; validation; visualization; writing—original draft; writing—review and editing.

RX, investigation; methodology; data curation; formal analysis; validation; writing—original draft; writing—review and editing.

DTZ, Methodology; data curation; writing—review and editing.

YM, Methodology; resource acquisition; data curation; writing—review and editing.

CS, Methodology; data curation; writing—review and editing.

SK, Methodology; data curation; writing—review and editing.

EK, Methodology; data curation; writing—review and editing.

PD, Methodology; data curation; writing—review and editing.

XQ, Methodology; data curation; writing—review and editing.

KAS, Methodology; data curation; formal analysis; resource acquisition; writing—review and editing.

MK, Methodology; data curation-—review and editing.

NM, Methodology; data curation; writing—original draft; writing—review and editing

MA, Resources, writing—review and editing

MAM, Methodology; data curation; writing—original draft; writing—review and editing

UW, Resources, writing—review and editing

CMC, Writing—review and editing

HI, Resources, writing—review and editing

RK, Methodology

CW, Methodology; review and editing.

KDCD, Conceptualization; writing—review and editing

KPN, Conceptualization; writing—review and editing

OOY, Resources; writing—review and editing

LM, Methodology; data curation; formal analysis; resources: validation; writing—original draft; writing—review and editing

BRR, Conceptualization; investigation; methodology; data curation; formal analysis; funding, acquisition; project administration; supervision; resources: validation; visualization; writing—original draft; writing—review and editing.

## ACKNOWLEDGEMENTS

The authors would like to thank Dr. Sarah Hill for providing cell lines and technical support. The graphical abstract was created using software from BioRender.com.

## Declaration of interest statement

H.I. is a consultant to Nanopath and Cellkey and receives research support from Canon USA, Nanoscope Systems, and ExoStemTech through sponsored research agreements through Mass General Brigham. These activities have no relationship to the study presented here.

All other authors have no conflicts of interest to disclose.

## Ethics approval statement

The Mass General Brigham Institutional Review Board approved this research.

## Patient consent statement

Patient-derived organoids were obtained under a secondary use protocol. No PHI was provided.

## Data availability statement

All data supporting the results have been included in the manuscript or as supplementary data.

## Funding statement

This work or the personnel were funded in part by Nile Albright Research Foundation (B.R.R. O.O.Y.), The Vincent Memorial Hospital Foundation (B.R.R.), NIH R37CA248565 (K.A.S.) NIH R01DK125263 (K.A.S.), Ovarian Cancer Research Alliance (K.A.S.); NIH R01GM138778 and R33CA281794 (H.I.); NCI–Cancer Center Support Grant P30 CA82709-25 Indiana University Simon Comprehensive Cancer Center (K.P.N.) and with gratitude to the Jerry and Peggy Throgmartin Family (K.P.N.); NCI 1U01CA284982 (C.M.C, B.R.R.) and R01CA264363 (C.M.C). NCI1P50CA240243 (C.M.C., B.R.R.).

## Permission to reproduce material from other sources

Not Applicable

## Clinical trial registration

Not applicable

## Notes

### Competing Interest Statement

H.Im is a consultant to Nanopath and Cellkey and receives research support from Canon USA, Nanoscope Systems, and ExoStemTech through sponsored research agreements through Mass General Brigham. These activities have no relationship to the study presented here. All other authors have no conflicts of interest to disclose.

