## Supplemental Material for "Extracellular vesicles facilitate the horizontal transfer of drug resistance and stem-like properties between ovarian tumor cells"

**Supplemental Data**

**Tables**

| **S.No.** | **EVs derived from** | **No. Particles (Particles/ml)** | **Size (nm)** |
| --- | --- | --- | --- |
| 1 | UWB-OlaSen cells | 2.5 × 10^11^ ± 2.30 × 10^11^ | 109.36 ± 16.25 |
| 2 | UWB-OlaRes cells | 4.46 × 10^11^ ± 9.27 × 10^11^ | 111.56 ± 4.43 |
| 3 | PEO-1 OlaSen cell | 3.8 × 10^11^ ± 4.78 × 10^11^ | 129.1 ± 1.7 |
| 4 | PEO1-OlaRes cells | 8.45 × 10^10^ ± 1.18 × 10^10^ | 133 ± 7.4 |
| 5 | 17-121 | 1.78 × 10^11^ ± 2.04 × 10^11^ | 126.96 ± 1.72 |
| 6 | VCRB330 | 0.66 × 10^10^ ± 0.63 × 10^10^ | 110.03 ± 6.89 |

**Table S1:** Particle number and size of extracellular vesicles (EVs) from olaparib sensitive and resistance cells and PDOs. The EVs were isolated from the olaparib drug sensitive and resistant cells and PDOs, and the particle number and size were determined by nanoparticle tracking analysis (NTA). Data represented as mean ± SEM (n = 3 independent experiments).

|  |  | **EV markers** | | | |
| --- | --- | --- | --- | --- | --- |
| **S.No.** | **EVs from** | **CD9** | **CD63** | **CD81** | **Calnexin** |
| 1 | UWB-OlaSen cells | 12.21 ± 2.88 | 60.52 ± 8.22 | 86.23 ± 8.47 | Negative |
| 2 | UWB-OlaRes cells | 19.6 ± 2.9 | 71.72 ± 11.1 | 95.86 ± 2.45 | Negative |
| 3 | PEO1-OlaSen cells | 38.9±5.91 | 20.1±21.34 | 71.8 ± 10.47 | Negative |
| 4 | PEO1-OlaRes cells | 97.1±10.23 | 52.9±10.89 | 99.3 ± 4.23 | Negative |
| 5 | 17-121 | 13.15 ± 0.22 | 42.1 ± 1.75 | 99.4 ± 0.15 | Negative |
| 6 | VCRB330 | 12.29 ± 1.98 | 50.93 ± 2.41 | 98.33 ± 0.45 | Negative |

**Table S2:** Relative expression of EV surface markers from olaparib sensitive and resistant cells and PDOs. The isolated EVs were subjected to flow cytometric analysis and evaluated with the EV markers CD9, CD63, and CD81. The negative marker of EV, calnexin was determined by immunoblotting. Data are represented as mean ± SEM (n = 3 independent experiments).

**Figures**

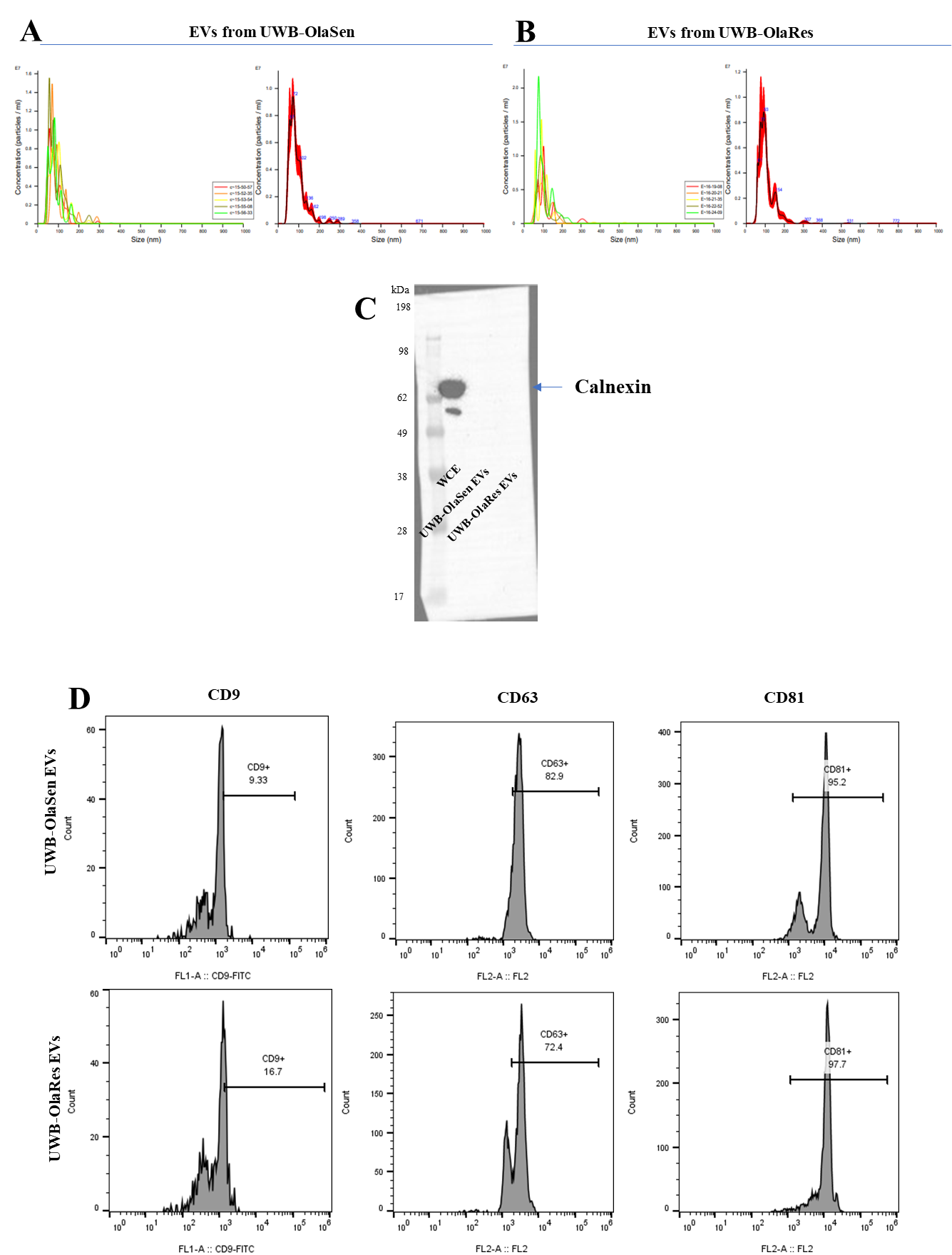

**Figure S1. Characterization of extracellular vesicles (EVs) from olaparib-sensitive and -resistant UWB1.289 mutant cells.** Particle size and concentration of EVs derived from (A) olaparib-sensitive (UWB-OlaSen) and (B) olaparib-resistant (UWB-OlaRes) UWB1.289 mutant cells were analyzed using NanoSight. (C) EV-negative marker, calnexin expression, was determined by western blotting in both EV preparations. (D) Surface expression of EV markers CD9, CD63, and CD81 was assessed by flow cytometry.

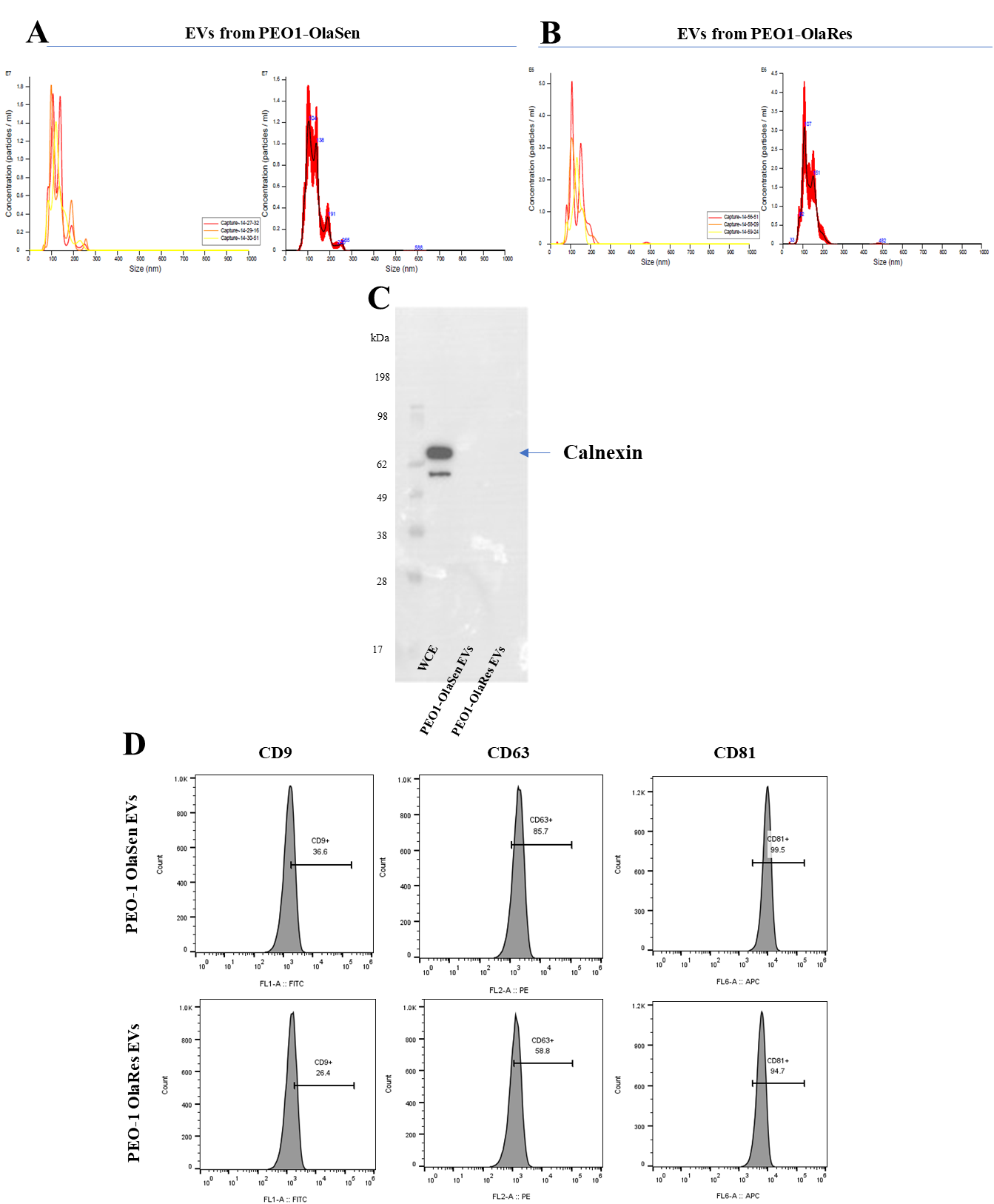

**Figure S2. Characterization of EVs from olaparib-sensitive and -resistant PEO1 cells.** Particle size distribution and concentration of EVs isolated from (A) olaparib-sensitive (PEO1-OlaSen) and (B) olaparib-resistant (PEO1-OlaRes) PEO1 cells, measured using NanoSight. (C) Calnexin expression was determined by western blotting in both EV preparations. (D) EV surface markers CD9, CD63, and CD81 in EVs derived from both olaparib-sensitive and -resistant cells analyzed by flow cytometry.

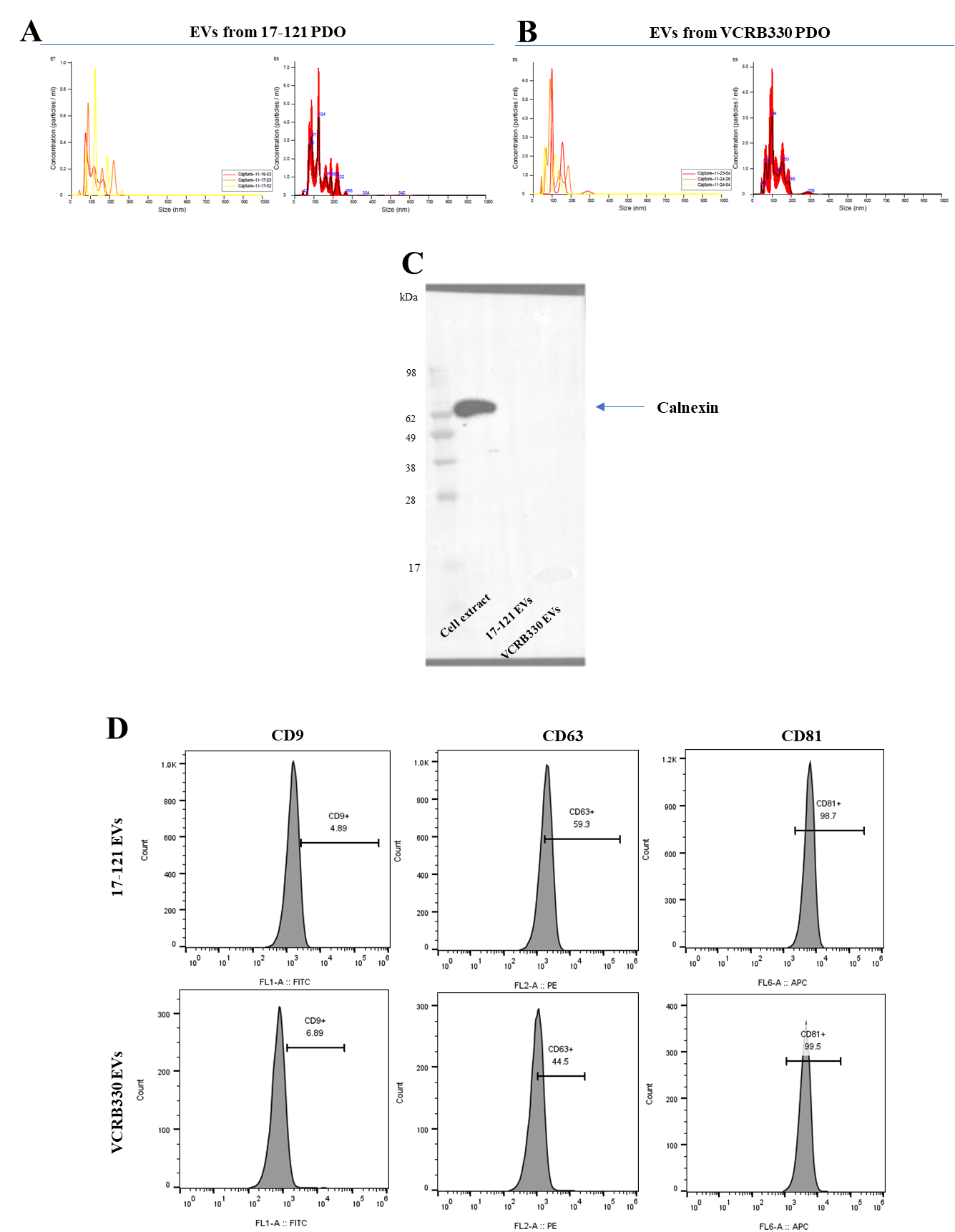

**Figure S3. Characterization of EVs from 17-121 and VCRB330 patient-derived organoids (PDOs).** Particle size distribution and concentration of EVs isolated from (A) 17-121 and (B) VCRB330 PDOs, measured using NanoSight. (C) Calnexin expression was determined by western blotting in both EV preparations. (D) EV surface markers CD9, CD63, and CD81 in EVs derived from both 17-121 and VCRB330 were analyzed by flow cytometry.

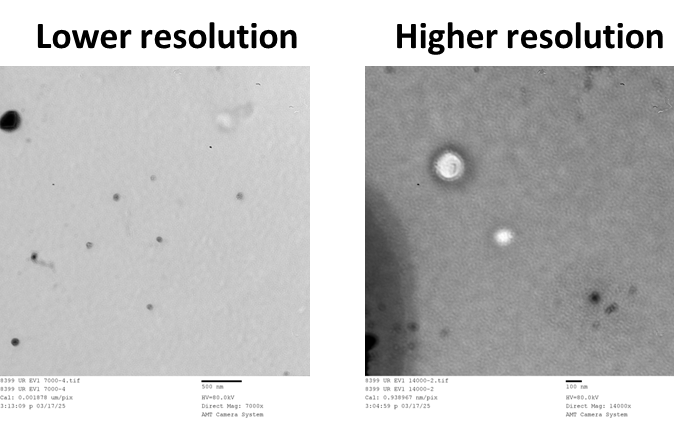

**Figure S4: Determination of extracellular vesicles (EVs) shape and size.** Transmission electron micrographs of extracellular vesicles isolated from UWB1.289 mut cells at bar=500 nm and bar=100 nm.

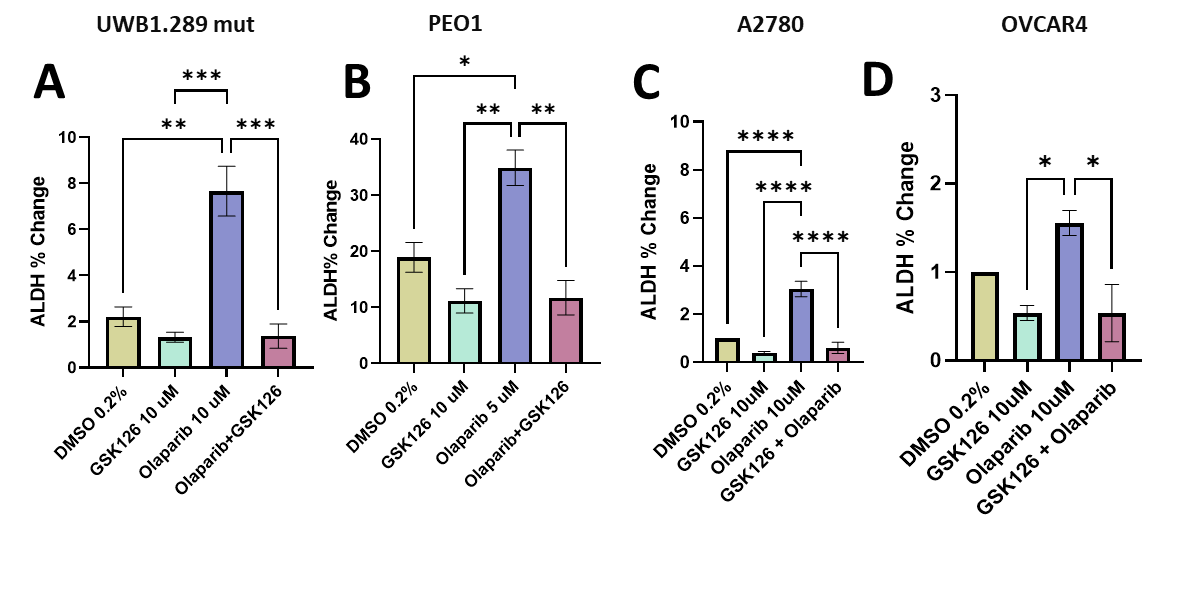

**Figure S5:** **Assessment of ALDH activity in human ovarian cancer cells following treatment with GSK126, olaparib, or their combination.** Flow cytometric analysis was used to measure ALDH activity in (A) UWB1.289 mut, (B) PEO1, (C) A2780, and (D) OVCAR3 cells. Significance was calculated using one-way ANOVA, with * indicating *P* < 0.05, ** indicating *P* < 0.01, *** indicating *P* < 0.001, and **** indicating *P* < 0.0001.

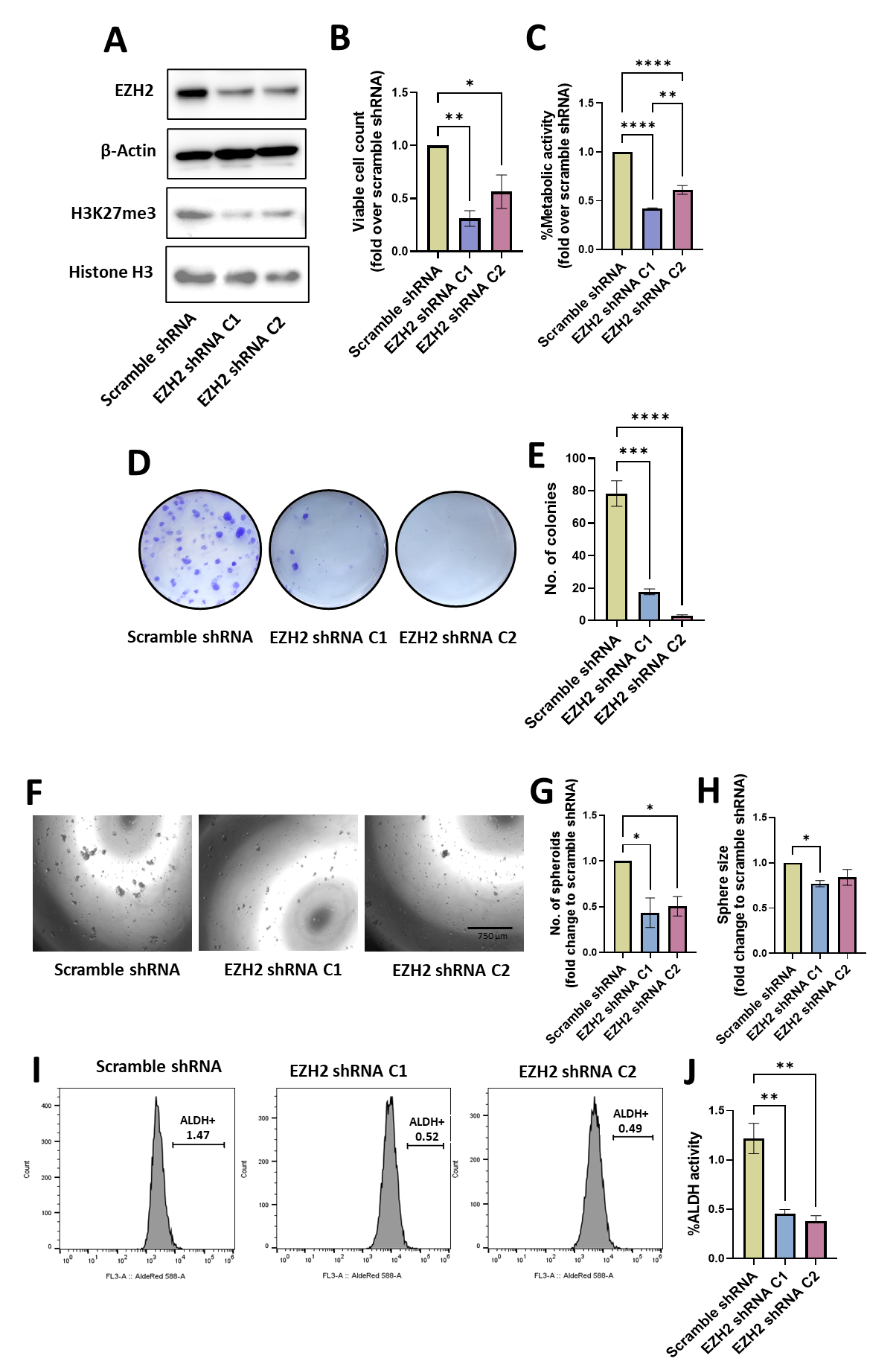

**Figure S6: Effects of EZH2 knockdown on cell viability, metabolic activity, clonogenic potential, sphere formation, and ALDH activity.** EZH2 was knocked down in UWB1.289 mut cells using the EZH2 Human shRNA Plasmid Kit. After puromycin selection, cells were sorted based on GFP expression into the following groups: scramble shRNA, EZH2 shRNA clone 1 (C1), and EZH2 shRNA clone 2 (C2). (A) EZH2 knockdown efficiency and H3K27me3 levels was confirmed by Western blotting (WB) in scramble shRNA and EZH2 shRNA-expressing cells. (B) Cell viability of scramble shRNA, EZH2 shRNA C1, and C2 was assessed using Toluidine blue staining and cell counts. (C) Metabolic activity was measured using the MTT assay. (D) Example of colony forming assay comparing the scrambled and EZH2 knockdown lines. (E) The number of colonies formed was quantified using ImageJ analysis. (F) Example of sphere-forming ability in scramble shRNA and EZH2 shRNA-expressing cells. (G, H) The number and size of spheroids were quantified. (H, I) ALDH activity in scrambled shRNA and EZH2 shRNA-expressing cells was assessed by flow cytometry. (J) Schematic representation of the effects of EZH2 knockdown in ovarian cancer (OvCa) cell lines. Significance was calculated using one-way ANOVA, with * indicating P < 0.05, ** indicating P < 0.01, *** indicating P < 0.001, and **** indicating P < 0.0001.

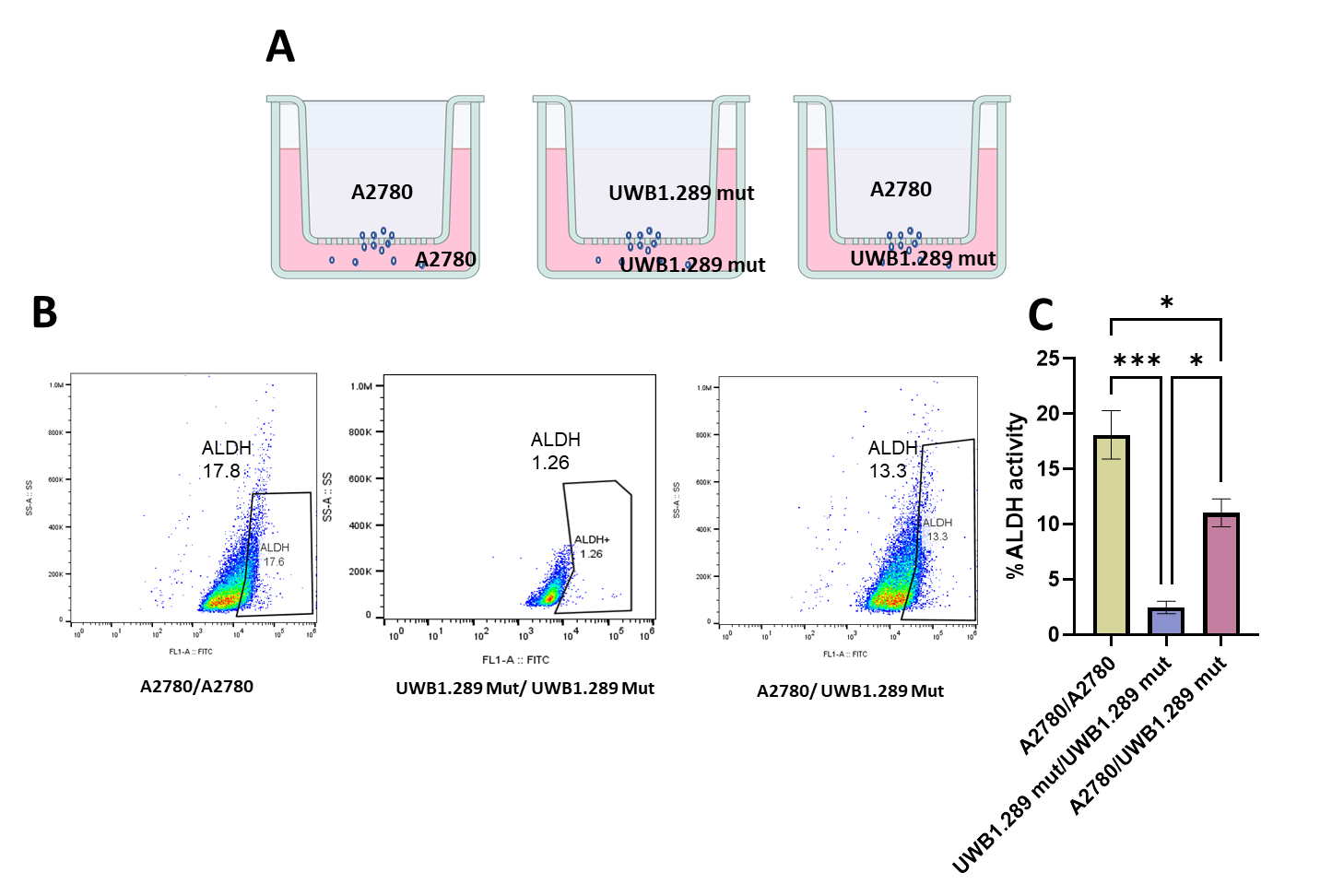

**Figure S7: Impact of cell-cell communication on ALDH activity in a co-culture system.** Schematic representation and quantification of ALDH activity in a transwell co-culture system. (A) Experimental setup: A2780 cells (high ALDH activity) were cultured in the upper chamber, while UWB1.289 mut cells (low ALDH activity) were placed in the lower chamber. Control conditions included A2780-A2780 and UWB1.289 mut-UWB1.289 mut co-cultures. A 0.4 μm porous membrane allowed exchange of soluble factors and small extracellular vesicles (EVs) while preventing direct cell contact. (B, C) Quantification of ALDH activity in UWB1.289 mut and A2780 cells under different co-culture conditions. Significance was calculated using one-way ANOVA, with * indicating *P* < 0.05 and *** indicating *P* < 0.001. Figure S3A was created in BioRender.com software.

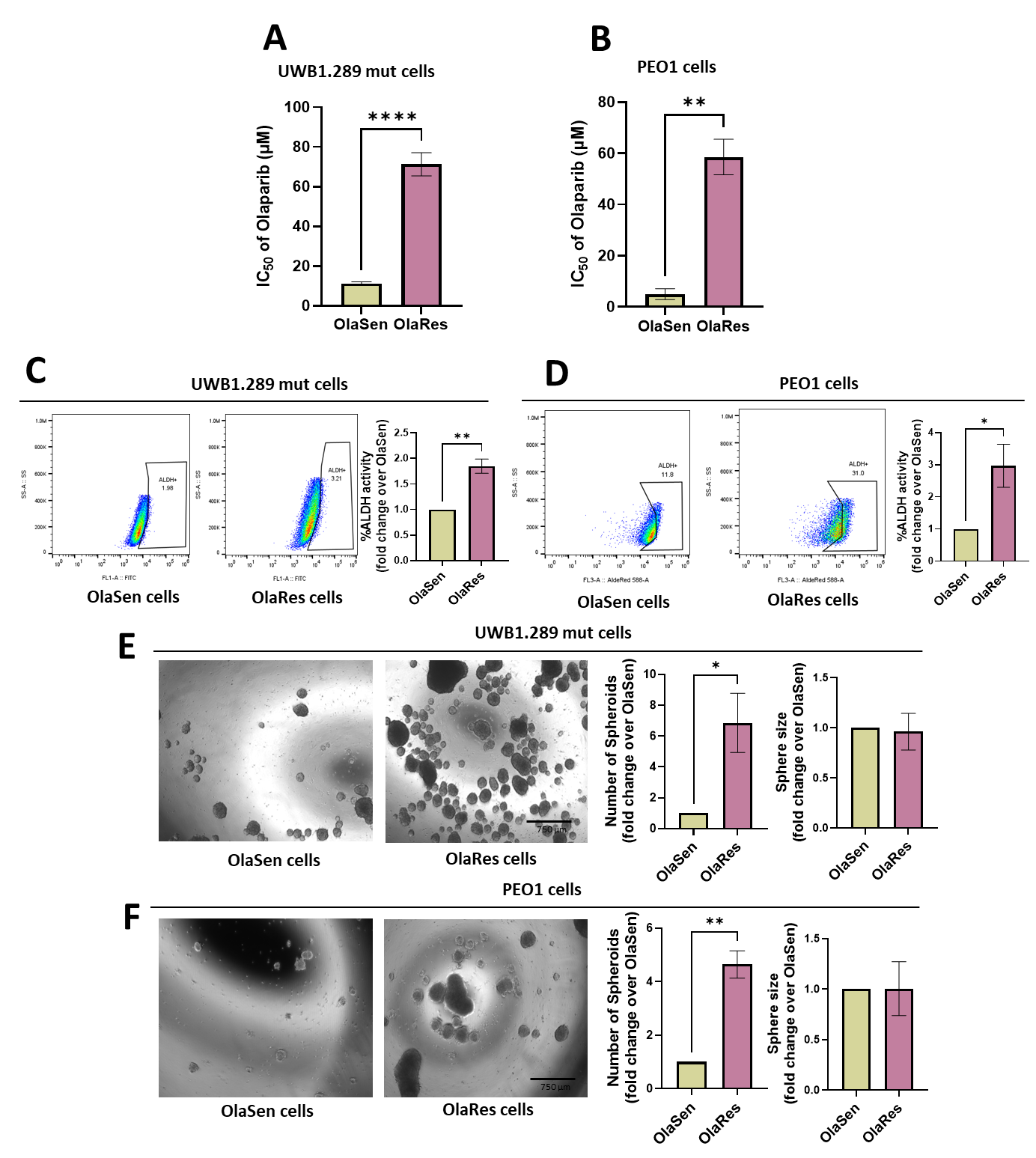

**Figure S8: Cell viability, ALDH activity, and sphere-forming ability in OlaSen and OlaRes cells.** Stable olaparib-resistant (OlaRes) cell lines were developed from (A) UWB1.289 mut and (B) PEO1 cells. The IC50 values of olaparib-sensitive (OlaSen) and olaparib-resistant (OlaRes) cells were determined using the MTT assay. Baseline ALDH activity of OlaSen and OlaRes cells was measured in (C) UWB1.289 mut and (D) PEO1 cell lines using flow cytometry. Sphere-forming ability was assessed in OlaSen and OlaRes cells of (E) UWB1.289 mut and (F) PEO1 lines. The number of spheroids and sphere sizes were quantified using ImageJ analysis. Significance was calculated using  *t*-test, with ** indicating *P* < 0.01 and **** indicating *P* < 0.0001.

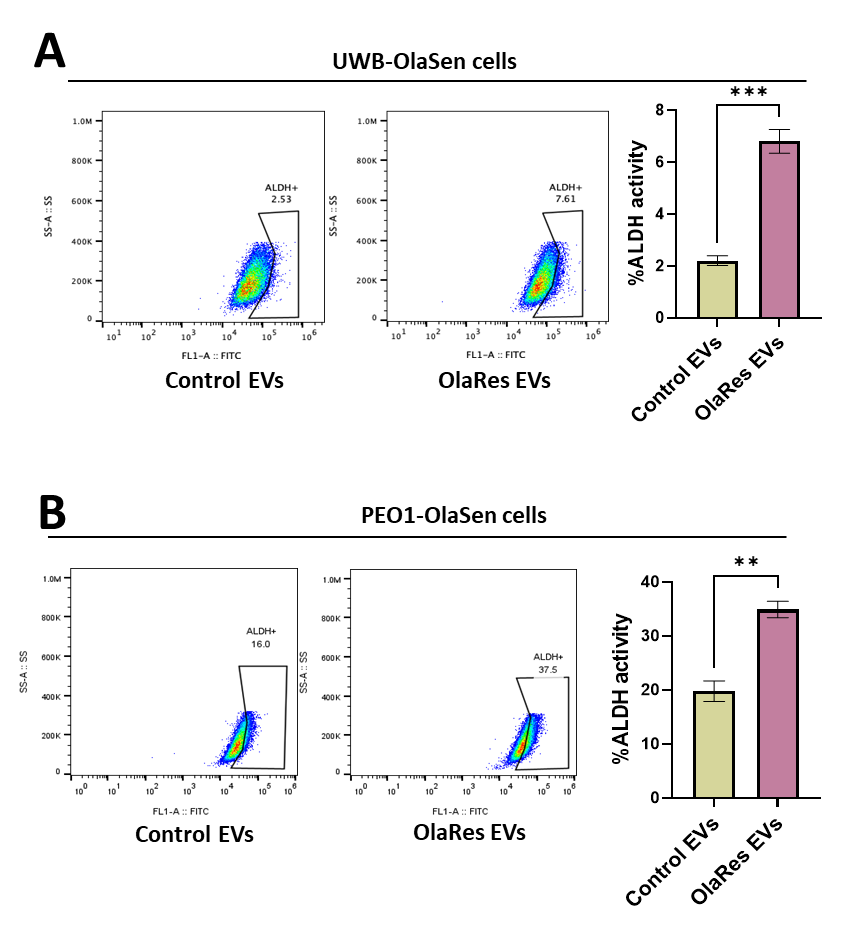

**Figure S9. Impact of EVs from olaparib-resistant cells on ALDH activity in drug sensitive cells.**

The effect of EVs derived from olaparib-resistant cells on ALDH activity was examined in (A) UWB1.289 mut olaparib-resistant (UWB-OlaRes) and (B) PEO1 olaparib-resistant (PEO1-OlaRes) cells, compared to EVs from control cells (derived from the corresponding parental olaparib-sensitive cell lines). Significance was calculated using *t*-test, with ** indicating *P* < 0.01 and *** indicating *P* < 0.001.

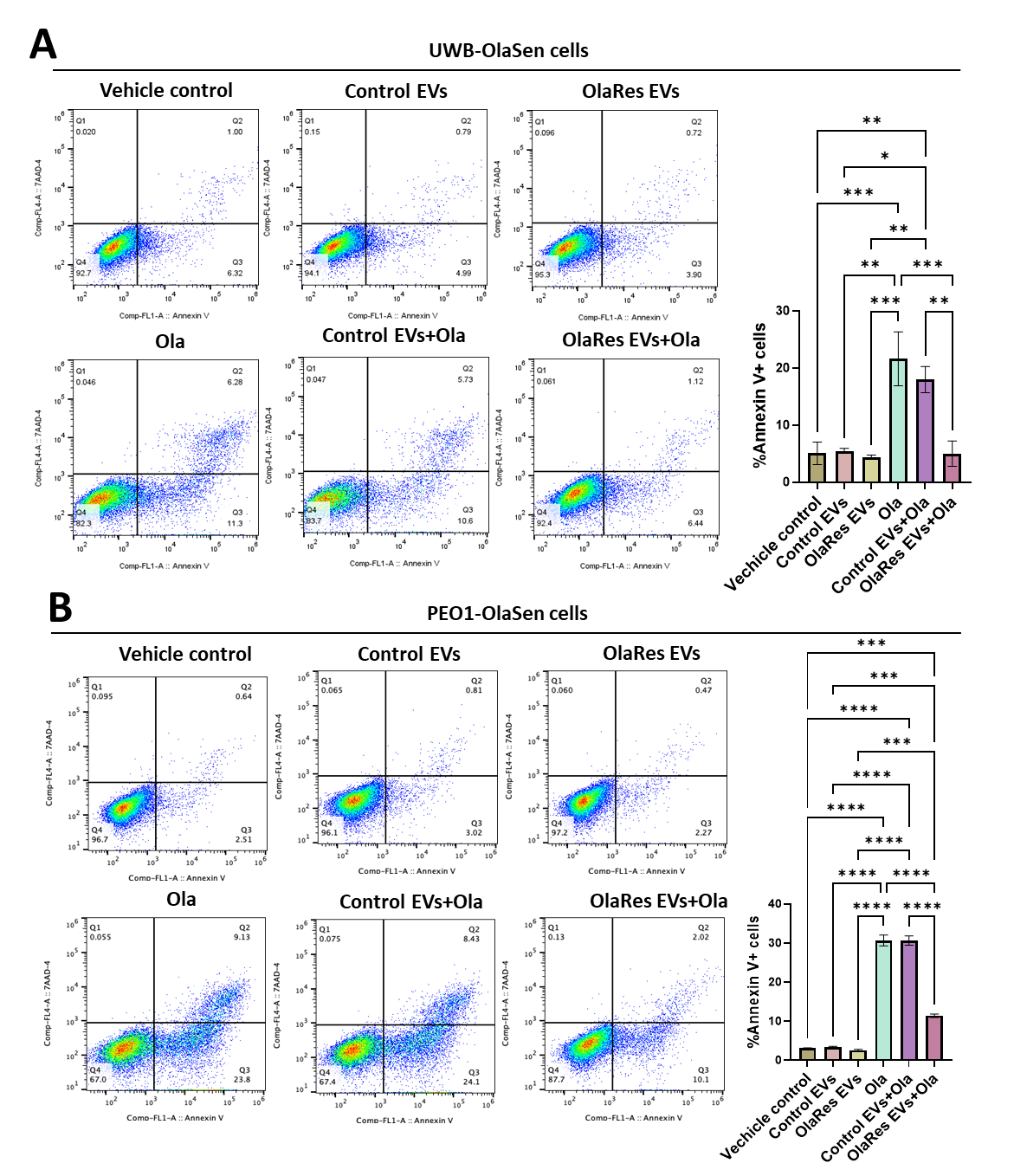

**Figure S10. Impact of EVs from olaparib-resistant cells on olaparib sensitivity and apoptotic response in drug sensitive cells.**

EVs were isolated from UWB-OlaRes and PEO1-OlaRes cells. UWB-OlaSen and PEO1-OlaSen parental cells were treated with EVs at a concentration of 2.5 µg/ml or vehicle and cultured for 24 hours (Day 0), prior to treatment with olaparib, with daily EV supplementation for 3 days. Apoptotic cell death in (A) UWB1.289 mut and (B) PEO1 parental cells was evaluated using annexin V binding assays. Significance was calculated using one-way ANOVA, with * indicating *P* < 0.05, ** indicating *P* < 0.01, *** indicating *P* < 0.001, and **** indicating *P* < 0.0001.

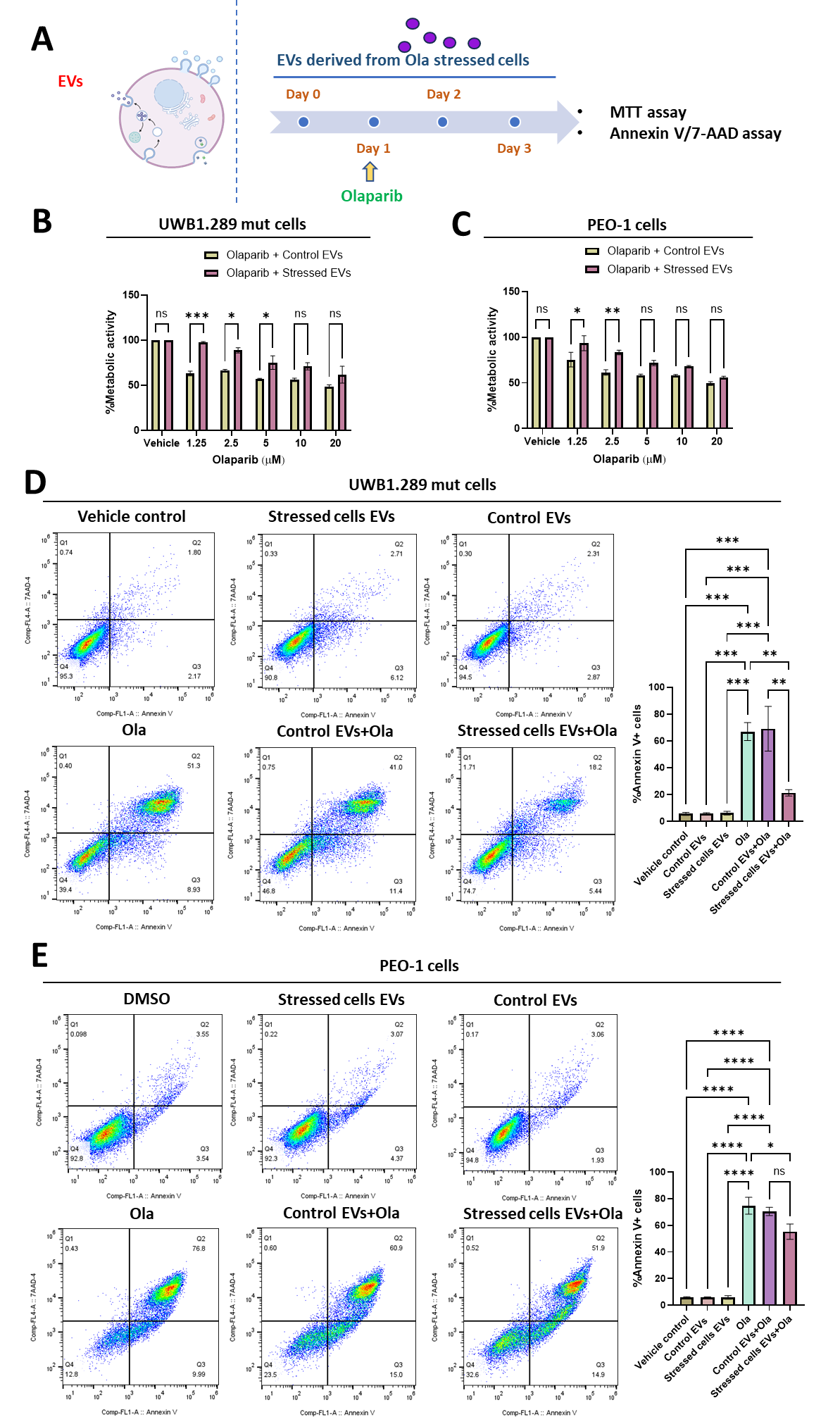

**Figure S11. Impact of EVs from acutely olaparib-treated cells on olaparib response in UWB1.289 mutant and PEO1 parental cells.** (A) EVs were isolated from UWB1.289 mut and PEO1 cells acutely treated with olaparib. UWB1.289 mutant and PEO1 parental cells were treated with EVs (2.5 µg/ml) or vehicle control and cultured for 24 hours (Day 0). On Day 1, cells were treated with 10 µM olaparib, followed by EV or vehicle supplementation on Day 1 and continued for an additional two days without further olaparib treatment. Metabolic activity of (B) UWB1.289 mutant and (C) PEO1 parental cells were assessed 72 hours post-treatment using the MTT assay. Apoptotic cell death in (D) UWB1.289 mutant and (E) PEO1 parental cells were evaluated using annexin V binding assays. Significance was calculated using one-way and two-way ANOVA, with * indicating *P* < 0.05, ** indicating *P* < 0.01, *** indicating *P* < 0.001, and **** indicating *P* < 0.0001. Figure S7A was created *in* BioRender.com software.
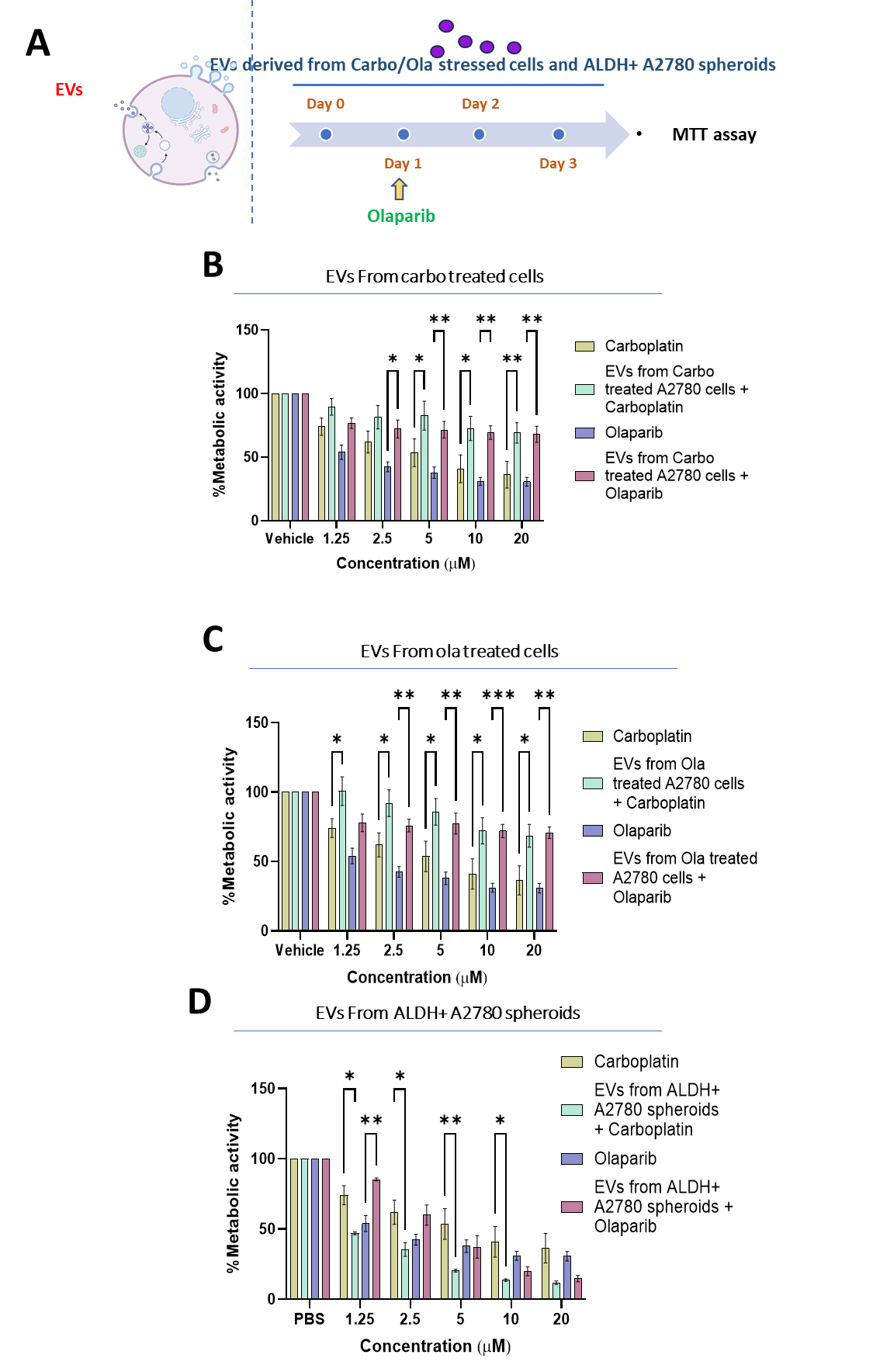

**Figure S12**: **Impact of EVs from carboplatin- and olaparib-treated A2780 cells and ALDH+ A2780 spheroids on metabolic activity of its parental cells.** (A) EVs were isolated from A2780 monolayer cells pre-treated with 5 µM of (B) carboplatin or (C) olaparib and collected 72 hours later, as well as from (D) ALDH+ enriched A2780 spheroids. Recipient cells were primed with EVs at a concentration of 2.5 µg/ml for 24 hours (Day 0) prior to treatment with carboplatin or olaparib, with daily EV supplementation for 3 days. Metabolic activity was assessed 72 hours post-treatment using an MTT assay. Significance was calculated using two-way ANOVA, with * indicating *P* < 0.05, ** indicating *P* < 0.01, and *** indicating *P* < 0.001. Figure S8A was created *in* BioRender.com software.

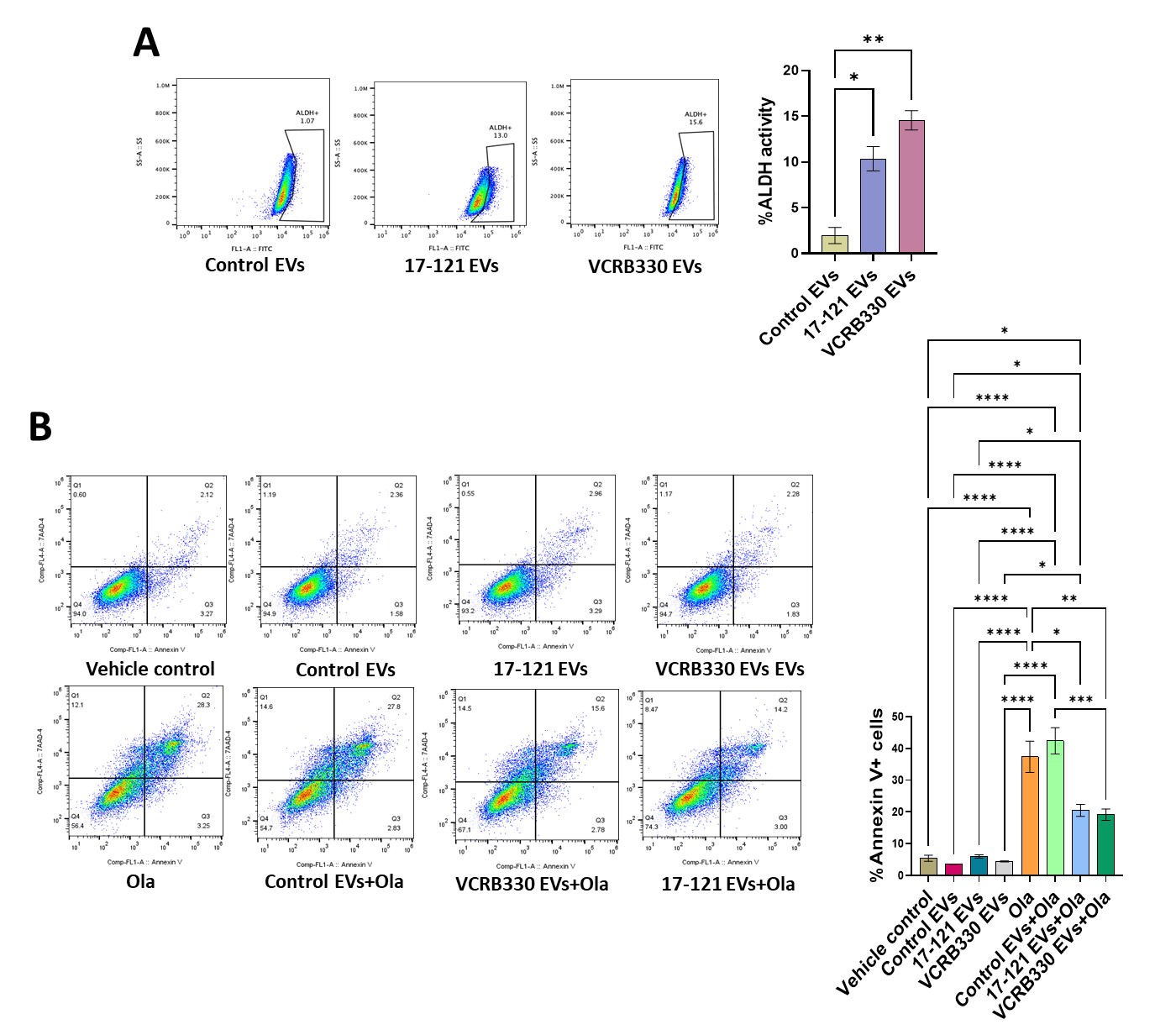

**Figure S13: EVs from HGSOC patient-derived organoids (PDOs) influence ALDH activity, olaparib sensitivity and apoptosis in UWB1.289 mut cells.** EVs were isolated from HGSOC PDOs 17-121 and VCRB330. (A) UWB1.289 mutant (mut) cells were treated with EVs from PDOs 17-121 and VCRB330 at a concentration of 2.5 µg/ml for 72 hours, followed by assessment of ALDH activity using flow cytometry. UWB1.289 mut cells were treated with EVs (2.5 µg/ml) or vehicle for 24 hours (Day 0), followed by additional EV supplementation on Day 1 and for two more days. Olaparib treatment was administered on Day 1 only. (B) Apoptotic cell death was assessed by annexin V binding assays. Statistical significance was determined using one-way and two-way ANOVA, with *P < 0.05, **P < 0.01, ***P < 0.001, and ****P < 0.0001.

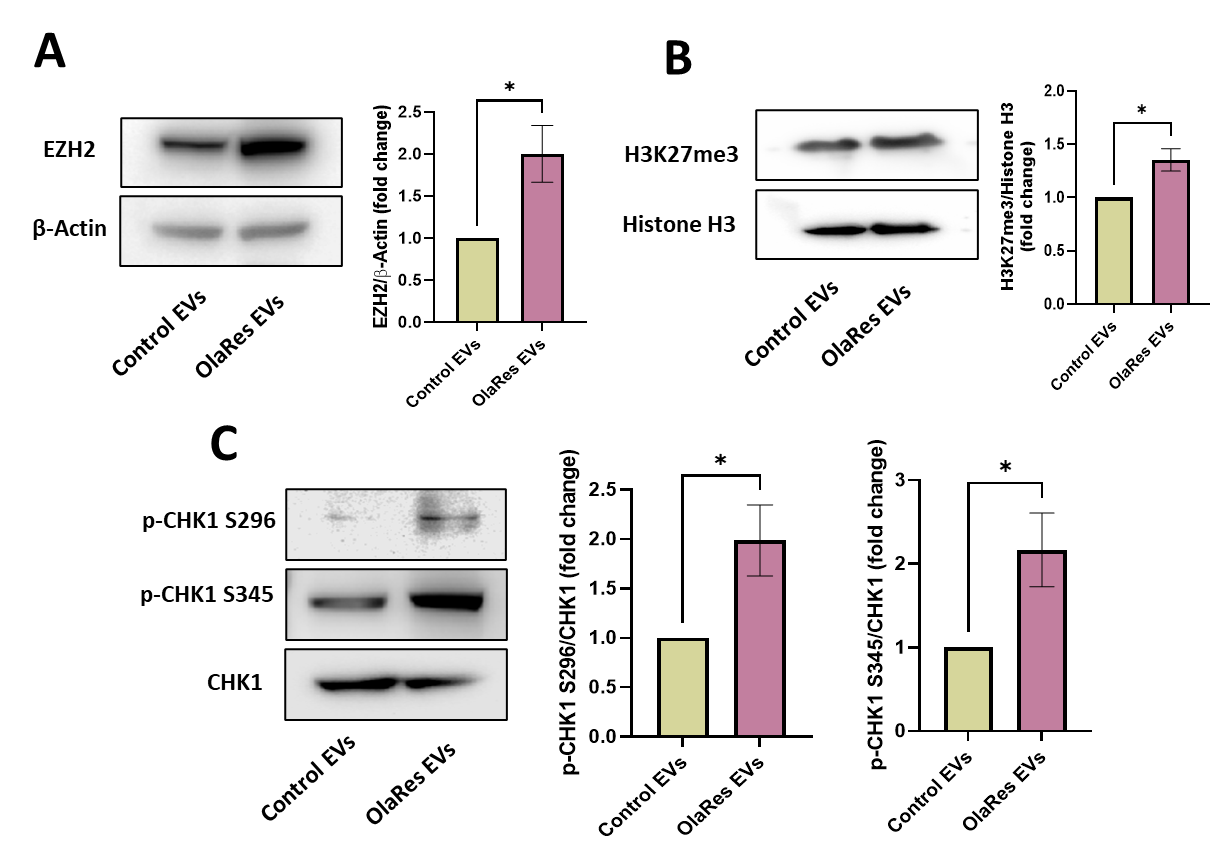

**Figure S14. Impact of EVs from PEO1-OlaRes cells on EZH2 expression, H3K27me3 levels, and CHK1 phosphorylation in its olaparib-sensitive cells.** PEO1-OlaSen cells were treated with EVs) drived from olaparib-resistant PEO1 (PEO1-OlaRes) cells or control cells. After 72 hours of incubation, western blot analysis was performed to assess (A) EZH2 and (B) H3K27me3 expression were analyzed in PEO1-OlaSen cells following EV treatment. (C) CHK1 phosphorylation at S296 and S345 was examined by western blotting after 30 minutes of EV incubation. Statistical significance was determined using *t*-test, with *P < 0.05.

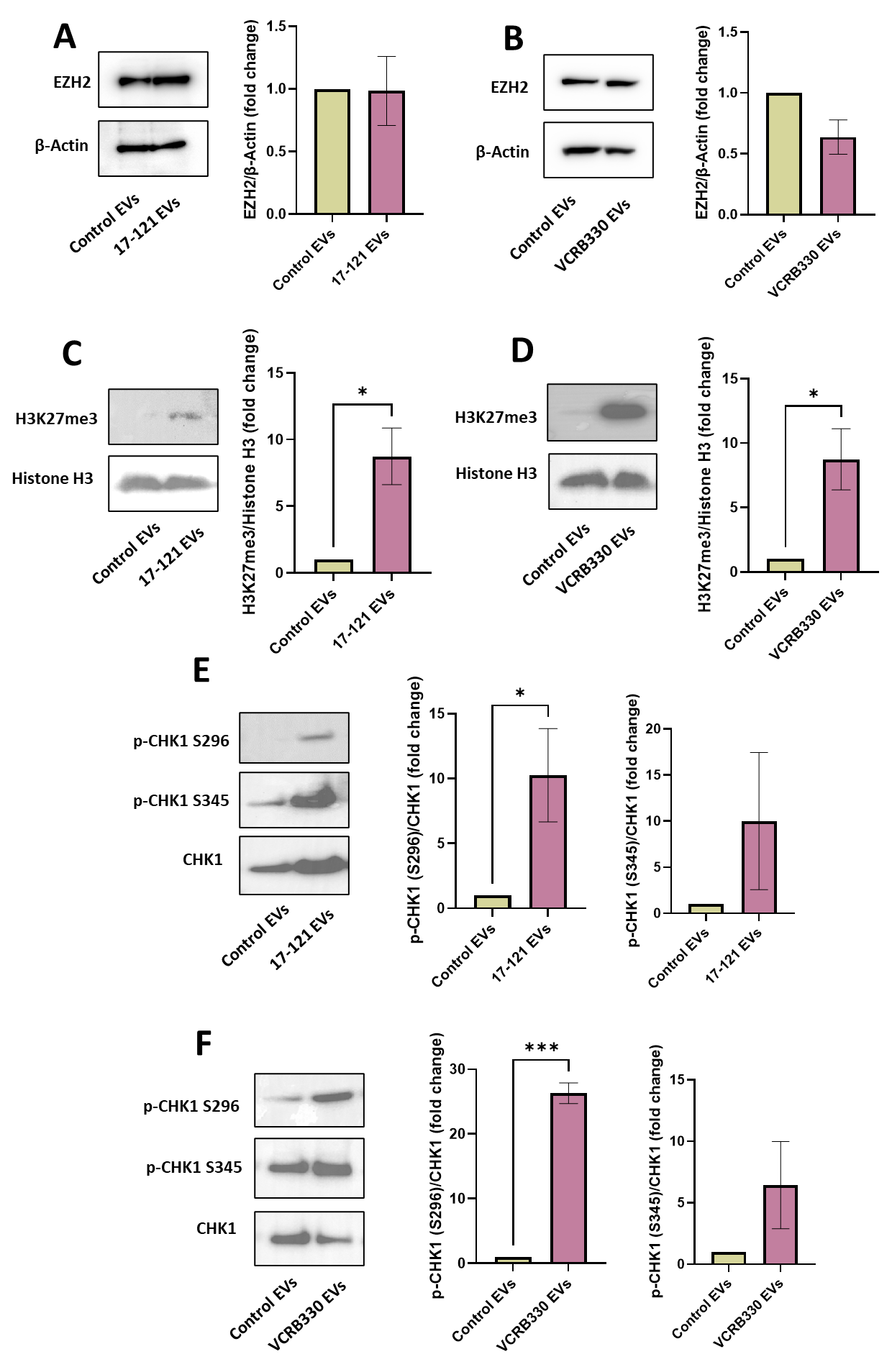

**Figure S15. Impact of EVs from 17-121 and VCRB330 PDO on EZH2 Expression, H3K27me3 levels, and CHK1 phosphorylation in BRCA1 mut cells.** UWB1.289 mut cells were treated with EVs derived from 17-121 and VCRB330 or control cells. After 72 hours of incubation, western blot analysis was performed to assess (A, B) EZH2 and (C, D) H3K27me3 expression were analyzed in UWB1.289 mut cells following EV treatment. (E, F) CHK1 phosphorylation at S296 and S345 was examined by western blotting after 30 minutes of EV incubation. Statistical significance was determined using *t*-test, with *P < 0.05 and ***P < 0.001.

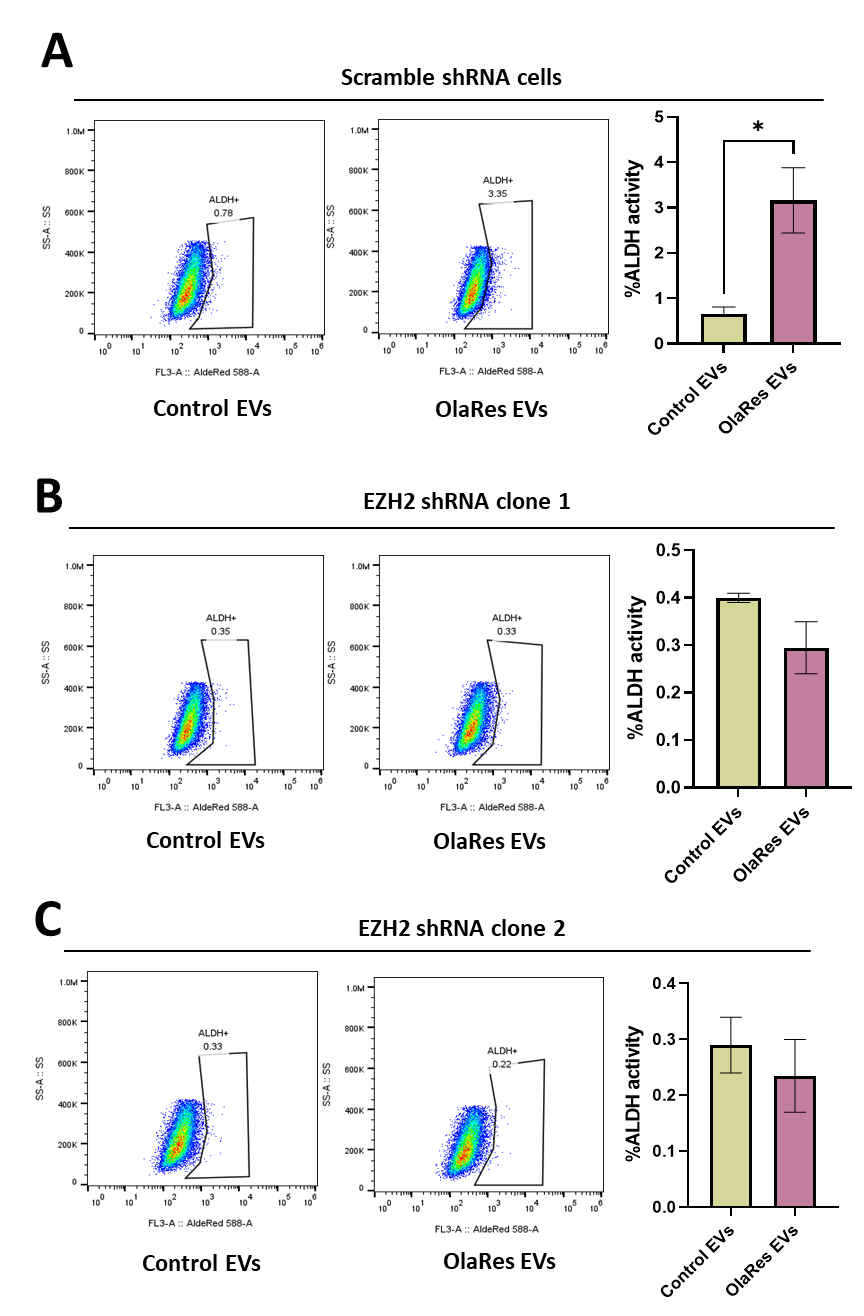

**Figure S16. Impact of EVs from olaparib-resistant cells stemness in scramble shRNA and EZH2 knockdown cells.** The effect of EVs from UWB-OlaRes or control cells on ALDH activity was evaluated using flow cytometry in (A) scramble shRNA , (B) EZH2 shRNA C1, and (C) EZH2 shRNA C2 cells. Statistical significance was determined using *t*-test and one-way ANOVA, with *P < 0.05.

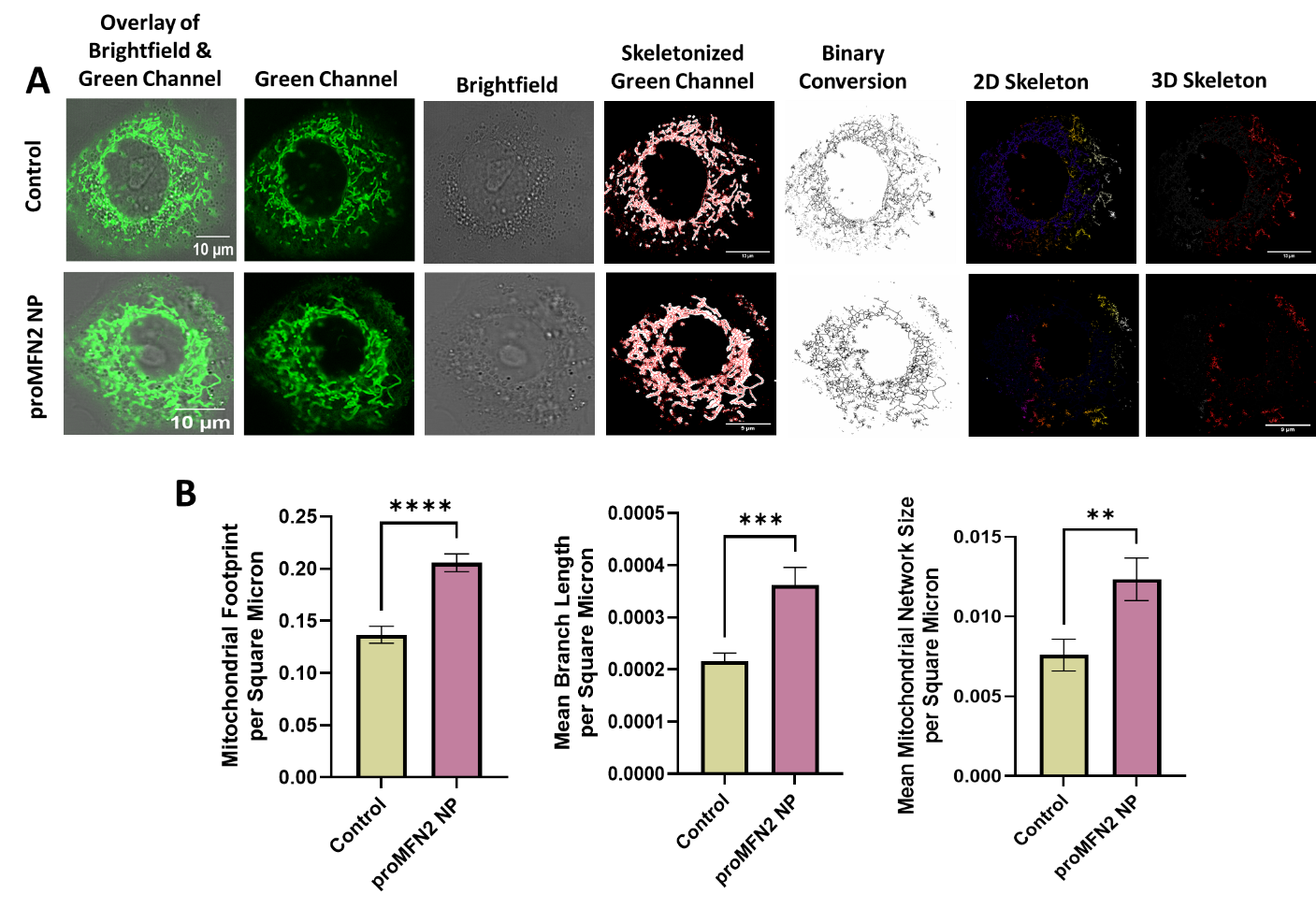

**Figure S17. Impact of proMFN2 NP on mitochondrial network remodeling in OvCa cells.** (A) Mitochondrial network analysis was performed by staining cells with MitoTracker Green after supplementing the cells with 10 µM proMFN2 peptide nanoparticles (NP), and as in Figure 7C, D, capturing images via confocal microscopy, and (B) quantifying mitochondrial parameters—including mitochondrial footprint, mean branch length, and mean mitochondrial network size—using ImageJ. Statistical significance was determined using a t-test, with and **P < 0.01, ***P < 0.001, and ****P < 0.0001.

**DETAIL METHODS**

1. **Cell lines and cell culture**

The UWB1.289 mut cell line carries a BRCA1 germline mutation (2594delC) in exon 11, along with a deletion of the wild-type allele. The PEO1 cell line harbors a homozygous BRCA2 mutation (5193C>G).

All the lines including A2780 and OVCAR4 OvCa were maintained under standard in vitro culture conditions in a humidified incubator at 37°C with 5% CO₂. The A2780 and OVCAR4 cells were cultured in RPMI-1640 medium (Cat #11,875,093, Gibco, Gaithersburg, MD), supplemented with 10% fetal bovine serum (FBS) (Cat #26,140,079, Thermo Fisher Scientific, Waltham, MA) and 1% penicillin-streptomycin (Pen-Strep) (Cat #15,070,063, Thermo Fisher Scientific). UWB1.289 mut cells were cultured in 1:1 mixture of RPMI-1640 and Mammary Epithelial Growth Medium (MEGM) (Cat #CC-3150, Lonza, Cambridge, MA), supplemented with 10% FBS and 1% Pen-Strep. PEO1 cells were cultured in RPMI-1640 medium supplemented with 10% FBS and 1% Pen-Strep. For the EV experiments, all cell lines were cultured and or treated in exosome-free serum medium (Cat #A2720801, Thermo Fisher Scientific). The cell lines were confirmed negative for mycoplasma prior to any assay by the MycoAlert Mycoplasma Detection Kit (Cat# LT07-318, Lonza).

2. **Establishment of human HGSOC organoid cultures**

Patient derived organoids were established as described earlier (1).

3. **Isolation of EVs**

EVs were isolated from conditioned cell culture medium using a sequential centrifugation, filtration, and ultracentrifugation protocol to ensure the removal of cellular debris and contaminants while preserving vesicle integrity. To remove large debris and cells from the conditioned medium, a series of low-speed centrifugation steps were performed. The cell culture medium was initially centrifuged at 200 × g for 5 minutes to eliminate large cellular debris. The supernatant was then subjected to 2,000 × g centrifugation for 10 minutes at 4°C to remove intact cells and apoptotic bodies. Following centrifugation, the clarified supernatant was further processed to remove smaller debris and concentrate EVs. The supernatant was filtered through a 0.22 µm sterile syringe filter (pre-wetted with Dulbecco’s Phosphate Buffered Saline (DPBS); Cat #14190144, Thermo Fisher Scientific) to eliminate any remaining cellular fragments and large EVs. The filtered supernatant was concentrated using a 100 kDa molecular weight cutoff (MWCO) centrifugal filter unit (Amicon® Ultra-15 Centrifugal Filter Unit; Cat #UFC910024, MilliporeSigma, St. Louis, MO). This step was performed at 3,000 × g for 30 minutes at 4°C, allowing for the retention of EVs while removing smaller soluble proteins and media components. The retentate was washed three times with sterile DPBS to eliminate any residual culture medium and ensure a pure EV preparation. To further purify and concentrate EVs, ultracentrifugation was performed using a Beckman Coulter Optima MAX-XP Ultracentrifuge equipped with a TLA-55 rotor (Cat #3574478, Beckman Coulter). The concentrated EV suspension was subjected to 100,000 × g (55K RPM) ultracentrifugation for 1.5 hours at 4°C to pellet EVs. A second round of ultracentrifugation at 100,000 × g was performed to further concentrate and purify the EVs by washing with DPBS. The EV pellet was resuspended in 200 µL sterile DPBS and aliquoted (~200 µL per sample). The EV aliquots were preserved at -80°C.

4. **EVs characterization by flow cytometry and western blot analysis**

EVs were characterized using flow cytometry with a bead-based capture assay and Western blot analysis for purity assessment. EVs were first bound to 2 µL of magnetic beads (Cat #A37304, Thermo Fisher Scientific) by incubating with 5 µL of EV sample and 75 µL of cell staining buffer at room temperature for 15 minutes with gentle mixing. A control sample without EVs was also prepared. To prevent non-specific antibody binding, 20 µL of Fc receptor (FcR) blocking reagent was added, followed by incubation on ice for 10 minutes. After washing with cell staining buffer and glycine treatment (100 mM in PBS) for 30 minutes, EV-bound beads were stained with FITC-CD9, PE-CD63, and APC-CD81 antibodies (BioLegend) at a final concentration of 1 µL per 100 µL and incubated at room temperature for 1 hour in the dark. The samples were then washed three times by centrifugation at 3,500 × g for 4 minutes to remove unbound antibodies before analysis by flow cytometry. To confirm the purity of the EV preparation, western blot analysis was performed to detect calnexin, an endoplasmic reticulum (ER) protein, using an anti-calnexin antibody (Cell Signaling Technologies). As calnexin is absent in EVs, its detection would indicate contamination from cellular debris. The absence of calnexin in the EV fraction validated the purity of the isolated EVs. Together, these assays confirmed the successful isolation and characterization of EVs based on surface marker expression (CD9, CD63, CD81) and the exclusion of intracellular contaminants.

5. **Characterization of EV size distribution and protein content**

The size distribution and concentration of the isolated EVs were determined using Nanoparticle Tracking Analysis (NTA) by Nanosight LM10 (Malvern, Framingham, MA). For optimal accuracy, the EV samples were first diluted in PBS to achieve an appropriate particle concentration that minimizes track overlap during analysis. Each sample was analyzed in triplicate to ensure reproducibility, with the corresponding NTA software tracking the Brownian motion of the particles to calculate both the average particle size and the total particle concentration. This approach provides a robust quantitative assessment of the EV preparations.

In addition to physical characterization, the protein concentration of the EV samples was determined using a microBCA assay kit (Cat# 23235, Thermo Fisher Scientific), following the manufacturer’s guidelines. This sensitive assay allows for the accurate quantification of total protein content in the EV preparations, which is critical for normalizing downstream applications and ensuring consistency across experiments. Together, these methods offer a comprehensive evaluation of the EVs by combining detailed information on their size, concentration, and protein content.

6. **Transmission Electron Microscopy (TEM)**

TEM was performed to assess the structure and size of EVs. EVs were fixed in a solution containing 2.5% glutaraldehyde and 2% paraformaldehyde, followed by TEM processing using a Philips CM-10 microscope (Eindhoven, The Netherlands), as previously described (2).

7. **Cell culture in exosome-free serum medium**

For the EV experiments, cells were cultured and treated using exosome-free serum medium (Cat #A2720801, Thermo Fisher Scientific) to ensure that any EVs detected originated solely from the cells under investigation. This specialized medium is formulated to be devoid of exogenous vesicles typically present in conventional serum supplements, thereby eliminating potential background noise and contamination that could compromise the accuracy of EV analyses.

8. **Spheroid formation assay**

To assess spheroid formation, cells were seeded into ultra-low attachment 6-well plates (Corning® Costar® Ultra-Low Attachment Plates; Catalog #7007, Corning, Glendale, AZ) at a density of 20,000 cells per well. The cells were cultured in a serum-free medium supplemented with essential growth factors, including epidermal growth factor (EGF), basic fibroblast growth factor (bFGF), and B-27™ supplement, to support spheroid formation. The cultures were maintained at 37°C in a humidified incubator with 5% CO₂, providing optimal conditions for spheroid growth. Spheroid formation was carefully monitored over a period of 10–14 days, with growth factors replenished every 3 days to sustain cellular viability and promote spheroid development. At the end of the incubation period, both spheroids and culture media were collected. The spheroids were allowed to settle, and the media was gently aspirated before resuspending the spheroids in a Phenol Red-free medium to eliminate background interference for downstream analyses. The number and size of the spheroids were quantified using ImageJ software, where spheroids were defined as distinct objects with a pixel size greater than 50 pixels. This quantification provided an objective assessment of spheroid formation efficiency and morphological characteristics in response to different treatment conditions.

9. **Establishment of A2780 ALDH+ spheroids**

A2780 cells were grown to confluence and then harvested for use in spheroid formation assays. Prior to seeding, the cells were labeled with an ALDH substrate using the ALDEFLUOR™ kit (Cat #01700, StemCell Technologies, Cambridge, MA) following the manufacturer's protocol, which enabled the identification of cells exhibiting high ALDH activity. These ALDH-positive cells were subsequently isolated via fluorescence-activated cell sorting (FACS). The sorted ALDH+ cells were then seeded at a density of 20,000 cells per well in ultra-low attachment 6-well plates using a serum-free medium enriched with growth factors to support spheroid development. The plates were incubated at 37°C in a humidified atmosphere with 5% CO2, and spheroid formation was monitored over a 10–14-day period.

10. **Impact of EVs from on sphere forming ability of drug-sensitive cells**

In this assay, olaparib-sensitive cells (UWB-OlaSen and PEO-OlaSes) are seeded at a defined density of 20,000 cells per well in ultra-low attachment 6-well plates. The cells are cultured in a serum-free medium supplemented with specific growth factors as described earlier. Here, EVs derived from either control or olaparib-resistant cells (UWB-OlaRes and PEO1-OlaSen) are added to the culture at the time of seeding and then replenished every 3 days along with the medium. Over a period of 13 days, spheroid formation is monitored, and the growth factors are replenished every 3 days to maintain optimal conditions. At the end of the incubation period, both the spheroids and the culture media are collected, allowed to settle, and resuspended in Phenol Red-free medium, which is used to enhance imaging clarity. Quantitative analysis of spheroid number and size is performed using ImageJ software, where spheroids are defined as objects exceeding 50 pixels in size.

11. **Generation of EZH2 Knockdown UWB 1.289 mut Cell Lines**

Lentiviral particles were produced by co-transfecting HEK293T cells with the EZH2 Human shRNA Plasmid (Cat #TL304713, OriGene, Rockville, MD) and necessary packaging plasmids. The packaging plasmids are essential to produce infectious lentiviral particles, which carry the specific shRNA construct designed to silence the EZH2 gene. After transfection, the viral supernatant, which contains the lentiviral particles, was collected from the culture medium. This supernatant was then filtered to remove any cellular debris and concentrated to increase the viral titer, ensuring efficient transduction of target cells. The viral particles were used to transduce UWB 1.289 cells. To facilitate the transduction process, 5 µg/mL polybrene was added to the culture medium. Polybrene acts as a transduction enhancer by increasing the efficiency of viral entry into the target cells. After 24 hours, the medium containing the lentiviral particles was replaced with fresh culture medium to remove excess virus and other reagents. Selection of successfully transduced cells was achieved using puromycin, an antibiotic that eliminates non-transduced cells. A concentration of 1 µg/mL puromycin was maintained for 7 days, allowing only the cells that had integrated the lentiviral construct (and hence, the resistance gene) to survive. After selection, the transduced cells were expanded and maintained in medium containing 1 µg/mL puromycin to ensure continued selection pressure. To identify and isolate the cells that were successfully transduced, GFP (green fluorescent protein)-positive cells were sorted using flow cytometry. GFP is expressed as a marker for successful transduction, allowing for easy identification and isolation of the cells that contain the shRNA construct targeting the EZH2 gene. The sorted GFP-positive cells were then cultured in medium containing 1 µg/mL puromycin to ensure continued selection. Finally, the efficiency of EZH2 knockdown was assessed using Western blot analysis.

Cells expressing scrambled, EZH2 shRNA clone 1, and clone 2 were plated in 10 cm dishes at a density of 0.5 × 10^6^ cells and incubated for 72 hours. After incubation, the cells were trypsinized, and protein concentrations were determined using the microBCA protein assay kit. Following normalization, SDS-PAGE analysis was performed as described in our previous work (3). A noticeable reduction in EZH2 protein expression in the transduced cells indicates effective gene knockdown, validating the success of the lentiviral shRNA system in silencing.

12. **Metabolic activity of EZH2 knockdown cells**

Cell metabolic activity was evaluated using the MTT assay. UWB1.289 mut scramble shRNA, EZH2 shRNA clone 1, and clone 2 cells were seeded in a 96-well plate at a density of 1,000 cells per well and allowed to adhere overnight. The cells were cultured for 72 hours. Subsequently, the medium was replaced with 100 µL of fresh medium containing 0.5 mg/mL MTT reagent (Cat # M6494, Thermo Fisher Scientific), and the cells were incubated at 37°C in a 5% CO2 atmosphere for 4 hours. After incubation, the supernatant was carefully removed, and the formazan crystals were solubilized in 200 µL of dimethyl sulfoxide (DMSO). Absorbance was measured at 570 nm using a microplate reader.

13. **Cell viability of EZH2 knockdown cells**

UWB1.289 mut scramble shRNA, EZh2 shRNA clone 1, and clone 2 cells were seeded at a density of 1 × 10^5^ cells per well in a 6-well plate and incubated under standard conditions for 72 hours. After the incubation period, the cells were trypsinized to detach them from the plate surface. The cell suspension was then stained with trypan blue, which selectively stains non-viable cells. The number of viable cells was determined by counting the cells using a Bio-Rad cell counter, which accurately differentiates between live and dead cells based on their ability to exclude the dye.

14. **Colony-Forming ability of EZH2 knockdown cells**

UWB1.289 mut scramble and EZH2 shRNA (Clone 1 and Clone 2) cells were plated at a density of 500 cells per well in 6-well plates. The cells were cultured under standard conditions and incubated for 14 days to allow colony formation. After the incubation period, the colonies were fixed with methanol and stained with crystal violet for 30 minutes. Images of the stained colonies were captured, and the number of colonies was quantified using ImageJ software for analysis.

15. **Spheroid forming ability of EZH2 knockdown cells**

UWB1.289 mut scramble and EZH2 shRNA (Clone 1 and Clone 2) cells were seeded into ultra-low attachment 6-well plates (Corning® Costar® Ultra-Low Attachment Plates; Catalog #7007, Corning, Glendale, AZ) at a density of 20,000 cells per well. The cells were cultured in a serum-free medium supplemented with essential growth factors, including epidermal growth factor (EGF), basic fibroblast growth factor (bFGF), and B-27™ supplement, to support spheroid formation. The cultures were maintained at 37°C in a humidified incubator with 5% CO₂, providing optimal conditions for spheroid growth. Spheroid formation was carefully monitored over a period of 13 days, with growth factors replenished every 3 days to sustain cellular viability and promote spheroid development. At the end of the incubation period, both spheroids and culture media were carefully collected. The spheroids were allowed to settle, and the media was gently aspirated before resuspending the spheroids in a Phenol Red-free medium to eliminate background interference for downstream analyses. The number and size of the spheroids were quantified using ImageJ software, where spheroids were defined as distinct objects with a pixel size greater than 50 pixels.

16. **ALDH activity of EZH2 knockdown cells**

ALDH activity in EZH2 knockdown cells was assessed using the ALDEFLUOR Kit. Briefly, UWB1.289 mut scramble and EZH2 shRNA (Clone 1 and Clone 2) cells were plated in 6-well plates at a density of 2.5 × 10^5^ cells per well and incubated for 72 hours. After the incubation, the cells were trypsinized, washed, counted, and resuspended in ALDEFLUOR buffer at a concentration of 2.5 × 10^5^ cells/mL. The cell suspension was divided into two aliquots, which were incubated with 5 µL of ALDEFLUOR substrate, either with or without 5 µL of the ALDH inhibitor DEAB. Following a 45-minute incubation at 37°C, the cells were washed with ALDEFLUOR buffer and analyzed using a FACS cytofluorimeter (Beckman Coulter Gallios, Brea, CA).

17. **Assessment of ALDH activity in HGSOC cell lines treated with EZH2 inhibitor GSK126, olaparib, and combination**

To evaluate the effect of the combination of the EZH2 inhibitor (GSK126) and olaparib on ALDH activity in HGSOC cell lines, UWB1.289 mut and PEO1, A2780, and OVCAR4 cells were plated in 6-well plates at a density of 0.1 × 10^6^ cells/well and allowed to attach overnight in complete growth medium. On Day 1, the cells were washed with PBS and treated with vehicle control (DMSO), GSK126, olaparib, or combination of GSK126 and olaparib in EVs-free serum media for 72 hours. On Day 4, the media was replaced with fresh EVs-free serum media, and the cells were re-treated with the respective compounds for an additional 72 hours, completing the treatment on Day 6. After treatment, the cells were washed with PBS, trypsinized, and resuspended in ALDH assay buffer from the ALDH Activity Assay Kit. The cells were then stained for ALDH activity following the manufacturer’s protocol, and analysis was conducted using flow cytometry to assess ALDH activity.

18. **Co-culture studies and ALDH activity assay**

Cells were co-cultured using 6-well transwell plates with a pore size of 400 nm to facilitate paracrine signaling between the high ALDH activity cells A2780 and low ALDH activity UWB1.289 mut cells while preventing direct cell-cell contact. In the experimental setup, A2780 cells (high ALDH activity) were cultured in the upper chamber, while UWB1.289 mut cells (low ALDH activity) were seeded in the lower chamber. Control conditions included A2780-A2780 and UWB1.289 mut-UWB1.289 mut co-cultures. The 0.4 μm porous membrane facilitated the exchange of soluble factors and small EVs between the chambers, while preventing direct cell contact. A total of 1 × 10⁶ A2780 cells were seeded in 1 mL of appropriate culture media in the upper chamber, and 0.5 × 10⁶ UWB1.289 mut cells were seeded in 2 mL of appropriate culture media in the lower chamber. ALDH activity was assessed at different time points (24, 48 and 72 hours). Consistent results were observed at the 72 hour timepoint, which was used for subsequent experiments. After the incubation, cells from the lower chamber were collected and analyzed for ALDH activity to evaluate their functional response to paracrine signaling.

19. **Development of olaparib-resistant cell lines**

olaparib-resistant UWB1.289 mut and PEO1 cell lines were generated through continuous exposure to progressively increasing concentrations of olaparib (Catalog #S1060, Selleckchem, Houston, TX) over a period of three months. The selection process began with a low concentration of 0.5 µM olaparib and was gradually escalated to a final concentration of 50 µM. Following the establishment of resistance, the olaparib-resistant cell populations were maintained in culture with a constant presence of 10 µM olaparib to ensure the persistence of the resistant phenotype.

20. **IC_50_ of OlaSen and OlaRes BRCA mutant OvCa cells**

To confirm the acquisition of resistance, OlaSen and OlaRes cells of UWB1.289 mut and PEO1 cell lines were plated in 96-well plates at a density of 1,000 cells per well. After 72 hours of incubation, metabolic activity was assessed using the MTT assay as described earlier. The half-maximal inhibitory concentration (IC_50_) values for the resistant cell lines were determined and compared to those of their respective drug-sensitive counterparts. Here, cells were exposed to DMSO as a vehicle control during the assay.

21. **ALDH activity of OlaSen and OlaRes BRCA mutant OvCa cells**

Baseline ALDH activity in OlaRes and OlaSen cells of UWB1.289 mut and PEO1 cell lines was assessed using the ALDEFLUOR Kit. Briefly, cells were plated in 6-well plates at a density of 2.5 × 10^5^ cells per well and incubated for 72 hours. After the incubation, the cells were trypsinized, washed, and counted. ALDH activity was then measured following the previously described.

22. **Sphere forming ability of OlaSen and OlaRes BRCA mutant OvCa cells**

Sphere-forming ability was assessed in OlaSen and OlaRes cells of UWB1.289 mut and PEO1 cell lines. Briefly, 20,000 cells were plated in ultra-low attachment 6-well plates and incubated for 13 days. The medium was replenished every three days. The remaining steps were performed as described previously.

23. **Cell-cell communication between OlaSen and OlaRes cells in a co-culture system**

Cell-cell communication between OlaSen and OlaRes cells of UWB1.289 mut and PEO1 OvCa lines was assessed using a transwell co-culture system. In this experimental setup, OlaRes cells (high ALDH activity) were cultured in the upper chamber, while OlaSen cells (low ALDH activity) were placed in the lower chamber. Control conditions included OlaRes-OlaRes and OlaSen-OlaSen co-cultures. A total of 1 × 10⁶ OlaRes cells were seeded in 1 mL of appropriate culture media in the upper chamber, and 0.5 × 10⁶ OlaSen cells were seeded in 2 mL of appropriate culture media in the lower chamber. The co-culture system was maintained for 72 hours under standard cell culture conditions. After the incubation, cells from the lower chamber were collected and analyzed for ALDH activity to evaluate their functional response to paracrine signaling.

24. **EV-mediated modulation of ALDH activity in drug-sensitive OvCa cells**

OlaSen UWB1.289 mut and PEO1 cells (2.5 × 10^5^ cells) were plated in 6-well plates and subsequently supplemented with EVs derived from OlaRes UWB1.289 mut and PEO1 cells, respectively. The cells were incubated for 72 hours, after which the ALDH activity of the drug-sensitive cells was determined as described previously.

25. **Impact of EVs derived from OlaRes cells on sphere forming ability of OlaSen cells**

UWB1.289 mut and PEO1 OlaSen cells were seeded into ultra-low attachment 6-well plates at a density of 20,000 cells per well. The cells were cultured in a spheroid growth medium. The cultures were maintained at 37°C in a humidified incubator with 5% CO₂, providing optimal conditions for spheroid growth. Spheroid formation was carefully monitored over a period of 13 days, with EVs from OlaRes UWB1.289 mut and PEO1 cells replenished along with medium every 3 days and followed as described previously.

26. **Impact of EVs from olaparib-resistant cells on olaparib sensitivity and apoptotic response in drug sensitive cells.**

UWB-OlaSen and PEO1-OlaSen parental cells were plated in 96-well plate with density of 1000 cells/well. Then the cells were supplemented with EVs from OlaRes cells and control cells (OlaSen cells) at a concentration of 2.5 µg/ml or vehicle and cultured for 24 hours (Day 0), prior to treatment with olaparib (10 uM), with daily EV supplementation for 3 days. Metabolic activity of UWB1.289 mut and PEO1 parental cells was assessed 72 hours post-treatment using an MTT assay as described previously. Similarly, apoptotic cell death in UWB1.289 mut and PEO1 parental cells was evaluated using annexin V binding assays, where cells were 2.5 × 10^5^ cells in 6-well plate. Here, DMSO employed as vehicle control. After 72 h, cells were trypsinized and washed and counted. The cell viability is determined by FITC-Annexin V/7-AAD (Cat# 640922, Biolegend) dual staining according to the manufacturer protocol.

27. **EVs from olaparib acutely treated cells on olaparib sensitivity and apoptotic response in its parental cells**

UWB1.289 mut and PEO1 cells were treated with 10 µM olaparib, while PEO1 cells received 5 µM olaparib. On Day 0, cells were seeded, followed by olaparib treatment on Day 1. A second dose of olaparib was administered on Day 4, and the experiment was concluded on Day 7. EVs were subsequently isolated from these acutely olaparib-stressed cells using the previously established protocol.

UWB1.289 mutant and PEO1 parental cells were seeded in a 6-well plate and allowed to adhere overnight. On Day 0, cells were treated with EVs derived from olaparib acutely treated UWB1.289 mut and PEO1-mut cells or control cells at a concentration of 2.5 µg/ml and cultured for 24 hours. On Day 1, cells were treated with 10 µM olaparib, followed by continued EV or vehicle supplementation for an additional two days without further olaparib treatment. 72 hours post-treatment, the metabolic activity of UWB1.289 mutant and PEO1 parental cells was assessed using the MTT assay following the manufacturer’s protocol. Absorbance was measured at 570 nm using a microplate reader to quantify metabolic activity.

Apoptotic cell death in UWB1.289 mutant and PEO1 parental cells was assessed using Annexin V binding assays. Cells were seeded in a 6-well plate and allowed to adhere overnight. On Day 0, cells were treated with EVs (2.5 µg/ml) or vehicle control and cultured for 24 hours. On Day 1, cells were treated with 10 µM olaparib, followed by continued EV or vehicle supplementation for an additional two days without further olaparib treatment. After 72 hours, cells were trypsinized, washed with PBS, and resuspended in Annexin V binding buffer. Apoptotic cell death was quantified using FITC-Annexin V/7-AAD dual staining according to the manufacturer’s protocol. Flow cytometric analysis was performed using a Gallios flow cytometer, and data were analyzed using FlowJo software.

28. **Impact of EVs from carboplatin- and olaparib-treated A2780 cells, as well as ALDH+ A2780 spheroids, on the metabolic activity of their parental cells**

EVs were isolated from A2780 monolayer cells treated with 5 µM of carboplatin or olaparib and collected 72 hours after treatment. Additionally, EVs were isolated from ALDH+ enriched A2780 spheroids as previously described.

A2780 parental cells were seeded at a density of 1,000 cells per well in a 96-well plate and primed with EVs at a concentration of 2.5 µg/ml for 24 hours (Day 0). Following priming, the cells were treated with either carboplatin or olaparib. Daily EV supplementation was continued for 3 days. Metabolic activity was measured 72 hours post-treatment using an MTT assay.

29. **Impact of EVs from HGSOC PDOs on ALDH activity**

EVs isolated from HGSOC PDOs 17-121 and VCRB330, as described earlier, were used to treat UWB1.289 mut cells. A total of 2.5 × 10⁵ cells were plated in 6-well plates and supplemented with 2.5 µg/ml of EVs derived from 17-121 and VCRB330 or control cells, respectively. After 72 hours of incubation, the ALDH activity of the drug-sensitive cells was assessed as previously described.

30. **Impact of EVs derived from HGSOC PDOs on olaparib sensitivity and apoptotic response drug sensitive cells**

UWB1.289 mut cells were seeded in a 96-well plate at a density of 1,000 cells per well. The cells were then supplemented with EVs derived from 17-121 and VCRB330, or control cells, at a concentration of 2.5 µg/ml and cultured for 24 hours (Day 0) before being treated with olaparib (10 µM). EV supplementation was continued daily for three days. Metabolic activity was assessed 72 hours post-treatment using an MTT assay, as previously described.

Similarly, apoptotic cell death in UWB1.289 mut parental cells was evaluated using annexin V binding assays. For this, 2.5 × 10⁵ cells were plated in a 6-well plate and treated with either EVs or vehicle (DMSO). After 72 hours, cells were trypsinized, washed, and counted. Cell viability was determined using FITC-Annexin V/7-AAD dual staining following the manufacturer’s protocol.

31. **Comet assay**

To investigate the impact of EVs derived from olaparib-resistant cells on the DNA damage response in drug-sensitive cells, both UWB1.289 mut and PEO1 OlaSen cells were seeded in 6-well plates at a density of 0.25 × 10⁶ cells per well. The following day, the cells were primed with EVs derived from olaparib-resistant or control cells at a concentration of 2.5 µg/ml. On the next day, the cells were treated with 10 µM olaparib or DMSO, in combination with the EVs. EV supplementation was replenished daily for a total incubation period of 72 hours. At the end of the incubation, cells were scraped, and 1 × 10⁵ cells were collected for the comet assay, according to the manufacturer's instructions (Comet Assay Kit, Cat #7100, Cell BioLabs, San Diego, CA). Briefly, cells were resuspended, embedded in agarose on microscope slides, and subjected to electrophoresis. DNA damage was evaluated by the formation of comet tails, indicating DNA strand breaks. Images were captured using a fluorescence microscope at 200× magnification.

32. **Impact EVs from OlaRes cells on EZH2 and H3K27me3 expression in OlaSen cells**

UWB-OlaSen and PEO1-OlaSen cells were individually seeded in 10‐cm dishes at a density of 5 × 10⁵ cells per dish. After allowing the cells to attach, they were treated with EVs isolated from UWB-OlaRes and PEO1-OlaRes cells, or with EVs from control OlaSen cells, at a concentration of 2.5 μg/mL. Following a 72-hour incubation period, the cells were harvested, lysed, and processed for Western blot analysis to evaluate EZH2 and H3K27me3 expression as previously described.

33. **Impact EVs from OlaRes cells on CHK1 phosphorylation in OlaSen cells**

Both UWB-OlaSen and PEO1-OlaSen cells were plated in separate 10‐cm dishes at a density of 1 × 10^6^cells per dish. Once the cells adhered, they were exposed to EVs derived from either UWB-OlaRes/PEO1-OlaRes cells or from control OlaSen cells, at a final concentration of 2.5 μg/mL. After a 30‐minute incubation period, cells were collected, lysed, and subjected to Western blot analysis to measure CHK1 phosphorylation at S296 and S345, as previously outlined.

34. **Evaluation of ALDH activity in UWB1.289 mut shRNA and EZH2 shRNA-modified cells following treatment with evs from UWB-OlaRes or control cells**

To assess the impact of EVs from UWB-OlaRes or control cells on ALDH activity, flow cytometry was performed on scramble UWB1.289 mut scramble shRNA, EZH2 shRNA C1, and EZH2 shRNA C2 cells. Briefly, cells were seeded in 6-well plates at a density of 1 × 10⁶ cells per well. The cells were treated with 2.5 μg/mL of EVs isolated from either UWB-OlaRes or control cells and incubated for 72 hours. ALDH activity was then measured using a specific ALDH substrate, and the data were analyzed by flow cytometry to quantify ALDH activity in the different cell lines (UWB1.289 mut scramble shRNA, EZH2 shRNA C1, and EZH2 shRNA C2).

35. **Western Blot analysis of EZH2 expression in scramble shRNA and EZH2 shRNA-modified cells treated with EVs from various sources**

For Western blot analysis of EZH2 expression, scramble shRNA, EZH2 shRNA C1, and EZH2 shRNA C2 cells were plated at a density of 0.5 × 10⁶ cells in 10-cm dishes. The cells were then treated with EVs derived from scramble shRNA, EZH2 shRNA C1, C2, UWB-OlaRes cells/control cells, at a final concentration of 2.5 μg/mL. After 72 hours of treatment, the cells were harvested, lysed, and equal amounts of protein were subjected to Western blot analysis and EZH2 expression was determined by normalizing with β-Actin as described earlier (4).

36. **Quantification of HOTAIR levels in OlaSen and OlaRes cells**

For RNA extraction from both UWB1.289 mut and PEO1 cell lines, 0.5 × 10⁶ cells were seeded in 10 cm dishes and cultured for 72 hours. Total RNA was isolated from cells using the sRNA extraction kit (Thermo Fisher) according to the manufacturer’s protocol. cDNA synthesis was carried out using the Maxima First Strand cDNA Synthesis Kit (Thermo Fisher). Quantitative PCR was performed using SsoAdvanced Universal SYBR Green Supermix (Bio-Rad) to assess HOTAIR expression. The following primers were used: HOTAIR forward 5′-CAGTGGGGAACTCTGACTCG-3′ and reverse 5′-GTGCCTGGTGCTCTCTTACC-3′, and GAPDH forward 5′-GTCAACGGATTTGGTCTGTATT-3′ and reverse 5′-AGTCTTCTGGGTGGCAGTGAT-3′. The PCR reactions were conducted on a CFX96™ Real-Time PCR System (Bio-Rad). HOTAIR expression levels in both cell lines were calculated using the ΔΔCt method, with normalization to GAPDH.

37. **HOTAIR expression in EVs isolated from OlaSen and OlaRes cells**

Similarly, EVs were isolated from OlaSen and OlaRes UWB1.289 mut and PEO1 cells as described earlier. RNA was extracted from the EVs, and HOTAIR expression levels were assessed by RT-PCR, as described in the previous section.

38. **Effect of EVs from OlaRes cells on HOTAIR levels in recipient cells**

Both OlaSen UWB1.289 mut and PEO1 cells were supplemented with EVs derived from OlaRes cells or control cells at a concentration of 2.5 μg/mL. The cells were incubated with the EVs for 6 hours. After the incubation, total RNA was extracted from the recipient cells, and HOTAIR expression levels were measured by RT-PCR. The expression was compared to baseline levels in cells which are supplemented with EVs from OlaSen cells, respectively.
